# Single–cell transcriptional and epigenetic mapping reveals cellular and molecular mechanisms driving non-ischemic cardiac fibrosis

**DOI:** 10.1101/2024.05.09.593315

**Authors:** Crisdion Krstevski, Gabriella E. Farrugia, Ian Hsu, Malathi S. I. Dona, Taylah L. Gaynor, Charles D. Cohen, Rebecca L. Harper, Thomas I. Harrison, Bethany Claridge, Auriane Drack, Patrick Lelliott, Helen Kiriazis, Aascha Brown, Julie R. McMullen, Daniel G. Donner, Sean Lal, David W. Greening, Alexander R. Pinto

## Abstract

Cardiac fibrosis is a major cause of cardiac dysfunction. Recently, single-cell genomic approaches have revealed in unprecedented resolution the orchestrated cellular responses driving cardiac fibrosis. Yet, the fibrosis-causing phenotypes that emerge in the heart following non-ischemic cardiac stress, and the transcriptional circuits that govern cell identity and drive fibrosis, are not well understood. Applying a paired multiomic approach, we reveal key transcriptional circuits, in mouse and human hearts, which are associated with fibrosis development following non-ischemic cardiac insults, independent of disease model, species or biological sex. Strikingly, we find the key regulatory events driving fibrosis are reversible at the single-cell transcriptional and epigenomic level, further pointing to key factors regulating fibrosis development and resolution. The transcriptional regulators identified in this study represent promising targets to ameliorate the development of fibrosis in the context of chronic stressors such as aging and hypertension.

## INTRODUCTION

Cardiac fibrosis—the excess deposition of extracellular matrix (ECM)—can lead to heart dysfunction and failure. Fibrosis accompanies many cardiovascular insults. These include ischemic insults such as myocardial infarction (MI), or in non-ischemic insults, such as systemic hypertension, where an expansion of interstitial and perivascular ECM occurs^1^. Determining the cellular and molecular drivers of fibrosis is critical for the development of clinical approaches for ameliorating the deleterious effects of fibrosis in different contexts.

We have only recently begun to understand with precision the diverse cell types that form the heart, including those that contribute to fibrosis. A catalyst for the rapid advance of this field has been the advent of single-cell genomics, which has enabled analysis of the cardiac cellulome—the integrated network of cells that form the heart—in high resolution^2–6^. This technology has made possible precise detection of cellular mediators of fibrosis^7–10^, including ‘activated’ fibroblast subtypes that emerge following non-ischemic insults in both humans and mice, and are characterized by high expression of ECM-related genes^9,10^. Single-cell genomics has also been applied to decipher the gene regulatory networks (GRNs) that drive fibrosis and fibroblast activation. Given they act as nodes of control, GRNs may theoretically be therapeutically manipulated to ameliorate fibrosis development. However, our understanding, in particular their plasticity once exposed to fibrogenic stimuli such as hypertension, is rudimentary.

Here we have applied a paired multiomic approach—where both transcriptomic and epigenomic data is extracted from individual nuclei—to analyze cells from human hearts, and those from mice undergoing angiotensin II (AngII)-induced hypertension and cardiac remodeling. This unique dataset allowed us to gain new insights into transcriptional regulation in the context of cardiac fibrosis. Our analysis identified key transcription factors (TFs) that drive cell identity and non-ischemic fibrosis irrespective of species, disease model or biological sex. Further, we found fibrosis resulting from hypertension is reversible with significant plasticity evident in key GRNs that drive cardiac fibrosis. This work offers a resource for understanding the GRNs that govern cardiac cell identity and provides important insights of the cellular mediators and transcriptional circuity that drive fibrosis and heart failure.

## RESULTS

### Paired single-nucleus RNA-seq and ATAC-seq for the analysis of mouse cardiac tissue

To determine the core GRNs and corresponding TFs governing non-ischemic cardiac fibrosis, we initially used a hypertension-induced cardiac fibrosis model. To achieve this, we infused both male and female C57BL/6J adult mice (12-14 weeks of age) with AngII (1.4 mg/kg/day) or saline (control) for 14 days. Additionally, we tested the tractability and reversibility of GRN changes elicited by AngII infusion, by analyzing hearts from mice that underwent 14 days of AngII infusion followed by a 14-day recovery period (AngII+Rec), where ectopic AngII infusion was halted (Figure 1A).

**Figure 1.**
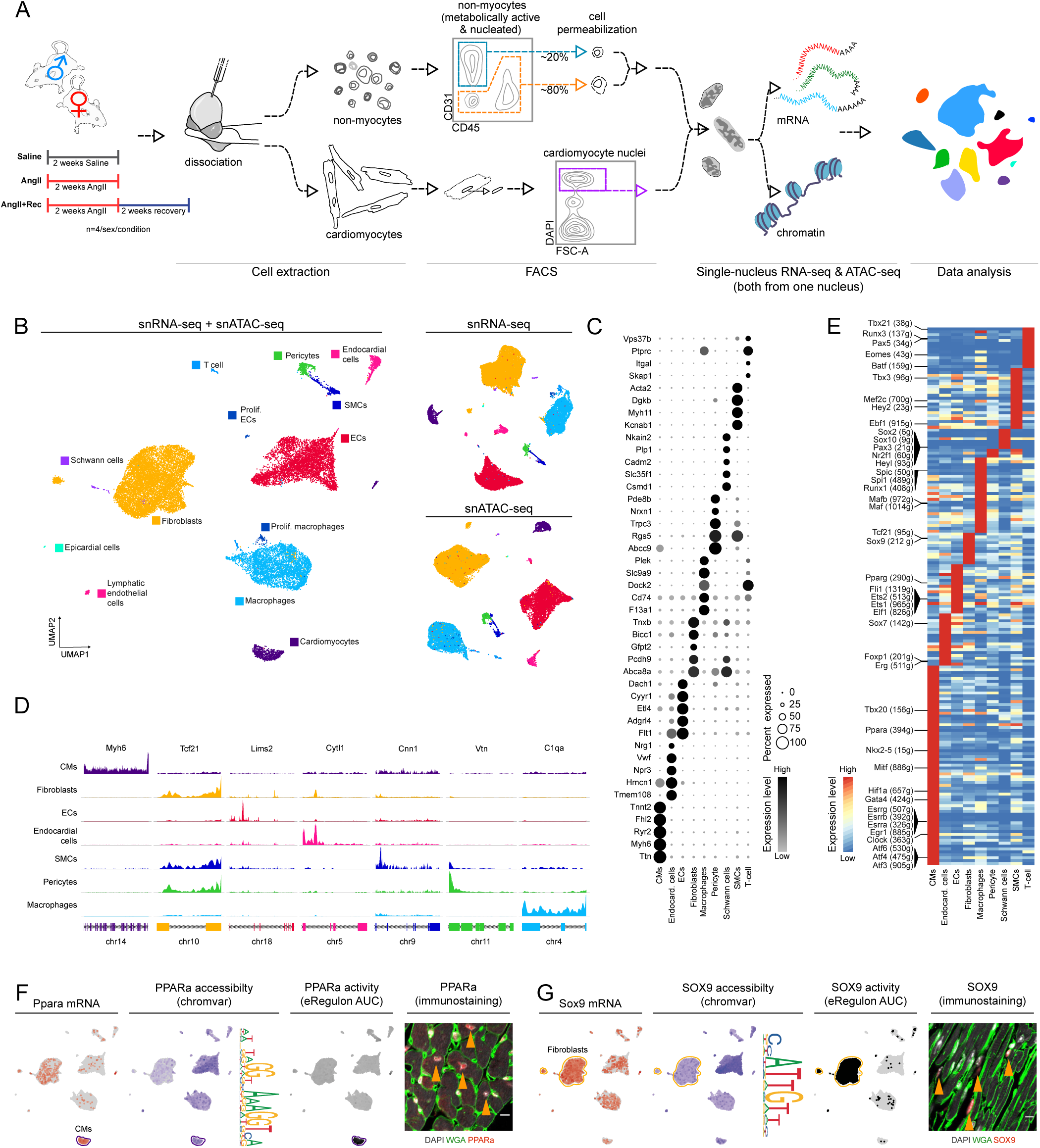
Paired single-cell transcriptomic and epigenomic profiling of the mouse heart. A. Experimental pipeline for performing paired single-nucleus RNA-sequencing (snRNA-seq) and single-nuclear Assay for Transposase Accessible Chromatin (ATAC)-sequencing (snATAC-seq) using mouse hearts with and without physiological stress. Left panel shows experiment cohorts used for mouse experiments: mice infused with saline or AngII for two weeks; or those infused with AngII for two weeks followed by an additional two weeks of recovery where ectopic AngII infusion was ceased. B. Uniform manifold approximation and projection (UMAP) of combined snRNA-seq and snATAC-seq data (left panel), snRNA-seq data alone (top right panel) or snATAC-seq data alone (bottom right panel). Each plot shows 22,182 cells. C. Top 5 distinct genes for each cell population determined by snRNA-seq. Dot size and intensity indicate cell proportion expressing the gene and relative gene expression level. D. Aggregate accessibility profiles for selected genes with relative cell-specific expression. Chromosome location and exons corresponding to genes indicated at the bottom. E. Enhancer-driven gene regulatory network (GRN) plot showing core eRegulons enriched within cardiac cell populations determined using the SCENIC+ pipeline. The heat map shows transcription factor (TF) expression of the eRegulon on the color scale. Number of TF target genes comprising the eRegulon indicated in parenthesis following the TF name. F. UMAPs corresponding to transcript level (grey-red, low-high, respectively), chromatin accessibility depicted using Chromvar package (grey-purple, low-high, respectively; with accompanying sequence logo) and eRegulon activity (grey-black, low-high, respectively) for Ppara. Accompanying micrograph shows Ppara in cardiac cells, with orange arrows indicating cardiomyocyte nuclei. Scale bar indicates 100 µm. G. Same as F, but for Sox9, and orange arrows indicating non-myocytes.

Before examining cardiac changes induced by AngII we characterized the ability of our system to understand gene regulators that drive fibrogenic phenotypes. We sought to develop a pipeline that considers both transcriptome and enhancer elements to identify GRNs that drive phenotypes of disparate cardiac cell types. Therefore, we performed parallel high-throughput profiling of transcriptomes and chromatin accessibility of individual nuclei of cardiac cells, using single-nucleus RNA sequencing (snRNA-seq) and single-nucleus Assay for Transposase-Accessible Chromatin sequencing (snATAC-seq) (Figure 1A, see Methods). Our paired analysis yielded a total of 22,182 nuclei that passed quality control measures (Supplementary Figure 1), where high-quality transcriptomic and genomic data could be retrieved, averaging a total of ∼7,231 nuclei per sample with a median number of 229,746 ATAC-seq fragments and 32,285 RNA-seq reads per nucleus. Dimensionality reduction and clustering of cells enabled classification of cell types based on paired transcriptomic and ATAC read outs (Figure 1B). We noted cell types could also be distinguished based on either transcriptomic (Figure 1B and 1C; Supplementary Figure 2A) or chromatin accessibility data alone (Figure 1B and 1D). Quantification of cells corresponding to individual timepoints showed relatively uniform contributions of cells from each sample (Supplementary Figure 1A-B).

Our dataset included all major cardiac cell types marked by distinct features. These include cell-type specific enrichment of transcripts, with those corresponding to key marker genes such as: *Ttn*, *Myh6* for CMs; *Dach1*, *Cyyr1*, *Flt1* for endothelial cells (ECs); *Npr3*, *Vwf* for endocardial cells; *Pcdh9*, *Tnxb* for fibroblasts; *F13a1, CD74* for macrophages; Abcc*9, Rgs5* for pericytes; *Myh11*, *Acta2* for smooth muscle cells (SMCs) (Figure 1C). Our dataset also captured extensive substructure within individual cell populations, with cell-specific transcriptional patterns within these cell subsets (Supplementary Figure 2B). Our snATAC-seq data showed cell-specific patterns of reads corresponding to genetic loci (Figure 1D). While open chromatin measures generally followed well-established patterns of cell-specific gene activity (Figure 1D), our findings complicate the capacity to use snATAC-seq alone to infer gene activity. For example, we note *Tcf21* gene locus is accessible in fibroblasts and mural cells (pericytes and SMCs; Figure 1D), despite *Tcf21* gene expression being restricted to cardiac fibroblasts^11^. This identifies a challenge for inferring gene activity using chromatin accessibility alone and highlights the value of accompanying paired transcriptomic data.

To determine GRNs underlying cell-type-specific phenotypes, we inferred transcriptional regulators using the SCENIC+ pipeline^12^. SCENIC+ leverages both epigenomic and transcriptomic data to infer enhancer-driven regulons (eRegulons)—genetic modules of TFs and their target genes. These are identified for each cell population using snATAC-seq data to determine enhancer accessibility and snRNA-seq data to determine TF and putative TF target gene expression. Correspondingly, this approach generates enhancer-driven GRNs (eGRNs). Using this pipeline, top distinct regulons identified for each cell type (Figure 1E) included those previously associated with specific cell types, such as Gata4, Ppara, Esrrg for CMs; Sox7, Foxp1 and Erg for endocardial cells; Fli1, and Ets1/2 for ECs; Sox9 and Pdgfra for fibroblasts; and Maf, Mafb, Spi1 and Runx1 for macrophages.

To verify cell-specific characteristics driving eRegulon discovery, we were able to assess features enabling cell-specific TF activity. Using Ppara and Sox9 (Figure 1F-G, respectively), for example, cell-type-specific eRegulon enrichment also accompanied enrichment of mRNA and accessibility of DNA binding sites for the respective TF. Further, immunostaining validated expression findings showing high PPARa (Figure 1F) in myocytes and SOX9 (Figure 1G, Supplementary Figure 5) in non-myocytes. However, our analyses also shows that neither TF gene expression nor the accessibility of the DNA binding sites for the TF alone are sufficient to yield transcription factor activity (Figure 1F-G). Therefore, while our pipeline enables discovery of TF activity, it also underscores the value of paired transcriptomic and epigenetic analysis for identifying TF activity governing cell identity.

### Paired single-nucleus RNA-seq and ATAC-seq analysis of human cardiac tissue

Next, we sought to determine if a similar paired multiomic approach could be applied to identify core GRNs driving diverse cellular phenotypes in the human heart. To achieve this, we mechanically dissociated cardiac samples from pre-mortem donor hearts from four females and four males with no clinical record of heart failure, extracted and sorted nuclei, and performed paired snRNA-seq and snATAC-seq (Figure 2A; see Methods). Our pipeline yielded 11,097 nuclei passing quality control criteria for transcriptomic and epigenomic data, representing major cardiac cell populations (Figure 2C; Supplementary Figure 3). Similar to our mouse dataset (Figure 1), human cardiac cell populations could be segregated based on snRNA-seq or snATAC-seq data. High enrichment of established cell-type-specific genes transcripts confirmed cell type identity (Figure 2D; Supplementary Figure 4). These included TTN, MYH6 for CMs, NEGR1 for fibroblasts; VWF for ECs, CD163, STAB1 for macrophages and MYH11 and CARMN for SMCs (Figure 2D). Again, we noted a substructure within individual cell types, most notably in CMs (Supplementary Figure 4B). Interestingly, while deconvolution of individual human samples contributing to the dataset revealed that non-myocyte cell subsets were composed of cells from all 8 individual donor samples, CM subsets mostly segregated based on individual donors (Supplementary Figure 4C). Examination of accessible chromatin also revealed cell-type specific patterns. These included MYH6 for CMs; DNMT1 for fibroblasts; FLT1 for ECs; CARMN for SMCs; PDGFRB for pericytes and C1QB for macrophages (Figure 2E). Finally, based on both snRNA-seq and snATAC-seq data, we identified eRegulons enriched within human cardiac cell types. These include GATA4 and PPARA in CMs, validated histologically (Supplementary Figure 5), and FLT1 in ECs, and SPI1 and EBF1 in macrophages (Figure 2F-E, Supplementary Table 1).

**Figure 2.**
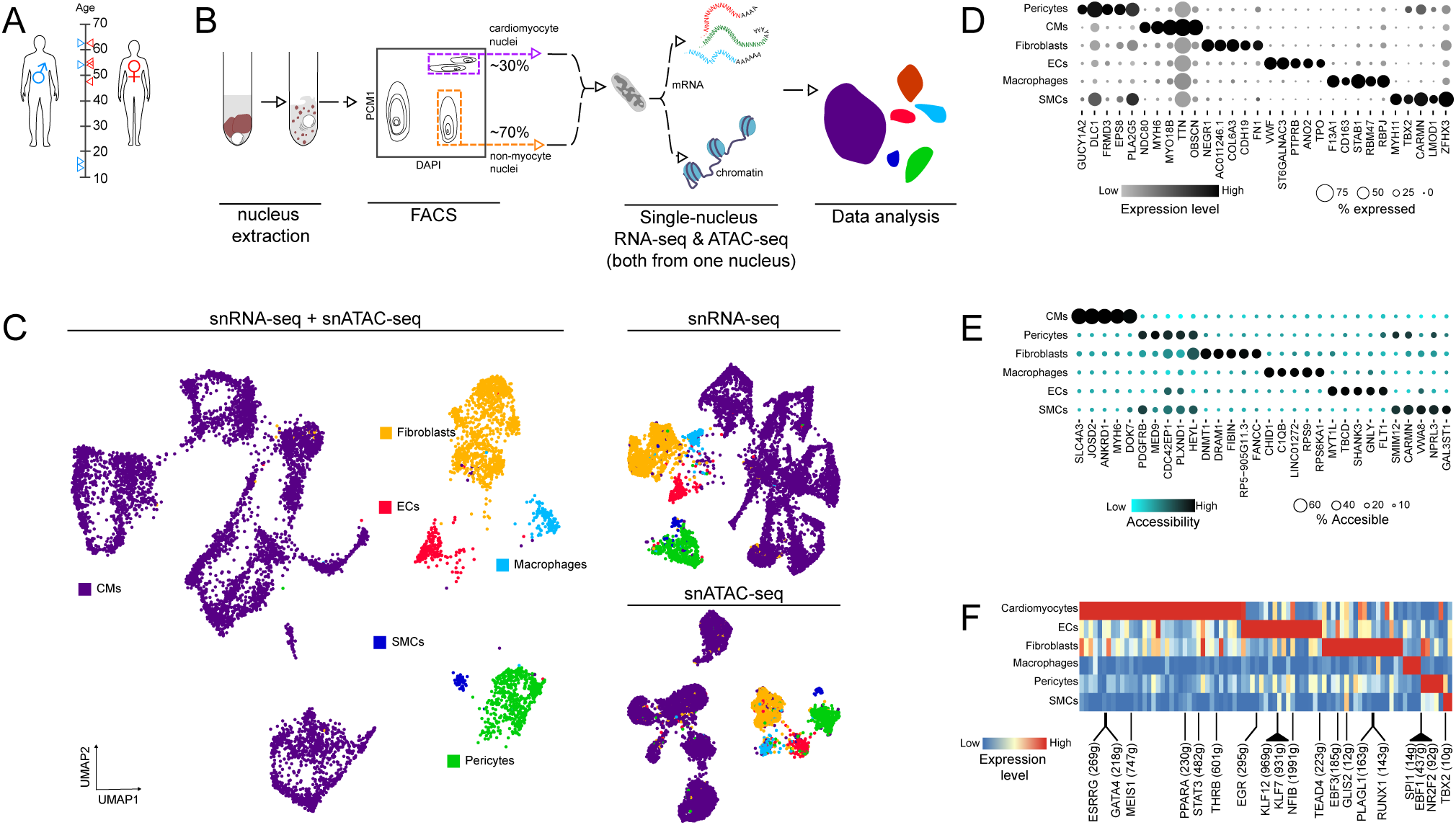
Paired single-cell transcriptomic and epigenomic profiling of human heart tissue. A. Gender, age and number of donor heart samples used for paired snRNA-seq and snATAC-seq. B. Pipeline for performing paired snRNA-seq and snATAC-seq following mechanical tissue dissociation of donor heart tissue. C. UMAP of combined snRNA-seq and snATAC-seq data (left panel), snRNA-seq data alone (top right panel) or snATAC-seq data alone (bottom right panel). Each plot shows 13,824 cells. D. Top 5 distinct genes for each cell population determined by snRNA-seq. Dot size and intensity indicate cell proportion expressing the gene and relative gene expression level. E. Genes associated with the top 5 accessible chromatin regions for each cell population determined by snATAC-seq. F. GRN plot showing core eRegulons enriched within cardiac cell populations determined using the SCENIC+ pipeline. The heat map shows TF expression of the eRegulon on the color scale. Number of TF target genes comprising the eRegulon indicated in parenthesis following the TF name.

Together, recovery of well-established master regulators of key cardiac cell types by our eGRN analysis of mouse and human hearts, demonstrate the advantages of applying a paired multiomic approach for discovering key transcriptional circuits governing cellular phenotypes.

### Multiomic analysis reveals gene regulatory network dynamics in non-ischemic fibrosis

Encouraged by our results, we applied this approach to determine core GRNs that drive development of cardiac fibrosis using the AngII-induced hypertension model. Picrosirius red (PSR) staining for collagen showed extensive cardiac fibrosis in response to hypertension in males and females (Figure 3A-B), with corresponding changes in a range of physiological parameters including systemic blood pressure (BP) (Supplementary Figure 6). Analysis of differentially expressed genes (p<0.01) between matching cell populations from Saline and AngII mice, showed significant changes in gene expression in cardiac cells (Figure 3C). Gene Ontology (GO) enrichment analysis mirrored cardiac fibrosis determined by histology, with genes induced by AngII corresponding to GO terms associated with ECM remodeling and transforming growth factor β (TGFβ) signaling, particularly by fibroblasts (Figure 3D). These included genes such as *Cilp*, *Col3a1* and *Jun*, associated with TGFβ response in fibroblasts, or *Postn*, *Col8a1, and Cnn2* associated with ECM remodeling (Supplementary Table 2). Among the top downregulated pathways in CMs were those related to cardiac contraction. These included genes such as *Gsn*, *Pde4d* and *Tpm1*. CMs also had several genes related to oxidative stress response downregulated, including *Map3k5*, *Cd36* and *Fos* (Figure 3D, Supplementary Table 2).

**Figure 3.**
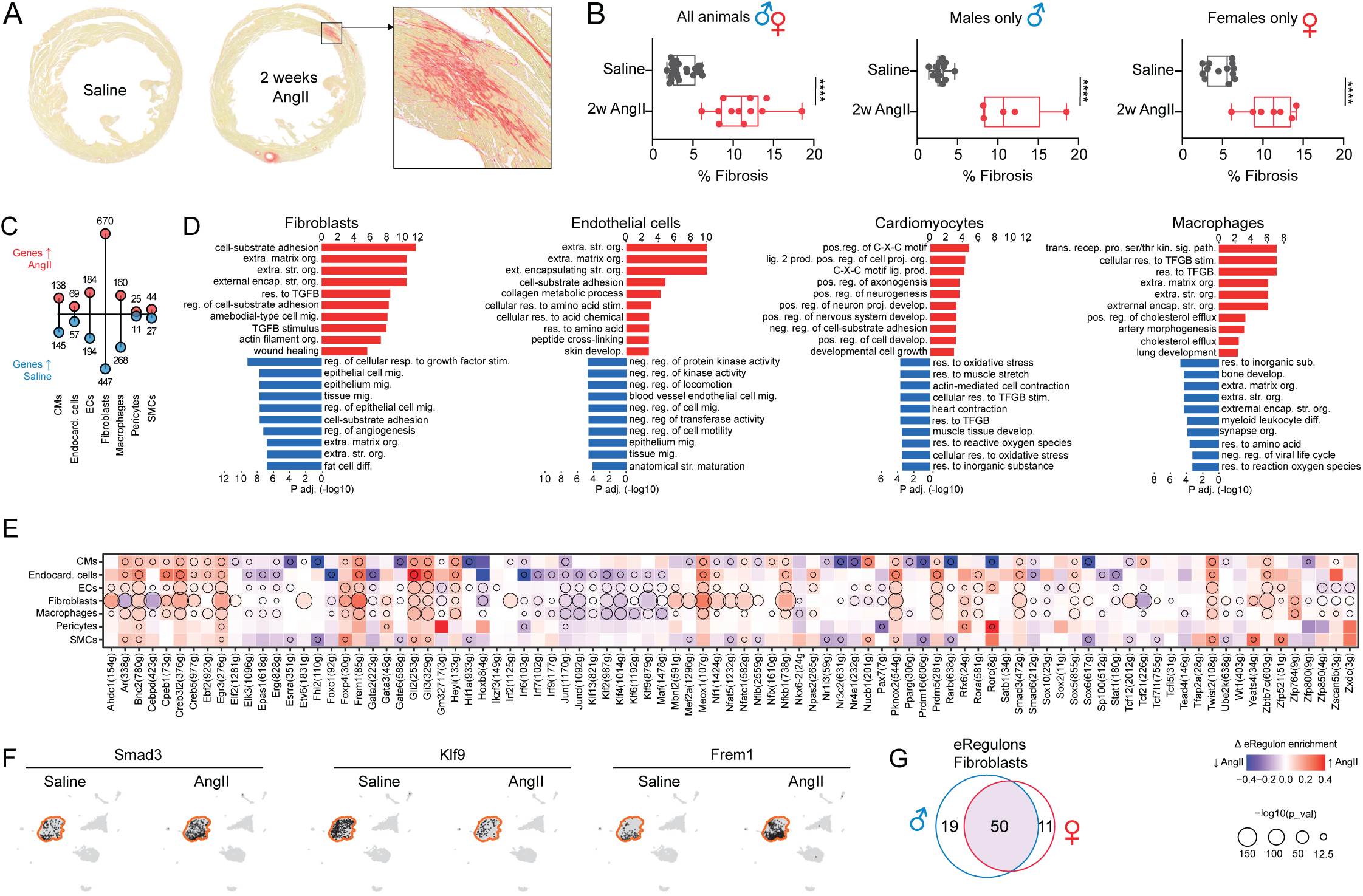
Transcriptional regulation of AngII-induced cardiac remodeling. A. Representative micrographs of picrosirius red stained cardiac cross sections from Saline or AngII mice. B. Change in cardiac fibrosis with and without induction of hypertension in both females and males (N= 5-6 animals per group). Cardiac fibrosis was measured from mid-ventricular sections. ****p <0.0001 (Wilcoxon test for differences in mean). C. Gene expression changes in major cardiac cell populations. Lollipop plot summarizing number of upregulated (red) and downregulated (blue) genes (p < 0.01) in AngII mouse heart cells relative to Saline heart cells. D. Gene Ontology (GO) enrichment analysis of genes that are up- and downregulated in fibroblasts, ECs, CMs and macrophages. E. Heatmap/dot plot summarizing changes in eRegulon enrichment in cardiac cells types between AngII and Saline mouse hearts. Heatmap color and intensity indicate direction and magnitude of change in eRegulon enrichment. Circle size indicates the statistical significance (Wilcoxon signed-rank test). F. UMAPs displaying binarized eRegulons enrichment within fibroblasts (outlined). Smad3 and Frem1 were chosen as example eRegulons that indicate activation of these TFs in AngII mouse cardiac fibroblasts. Klf9 was chosen as an example of a TF that is repressed in cardiac fibroblasts after AngII treatment. G. Venn diagram of eRegulons that are more highly enriched in cardiac fibroblasts after AngII treatment in males and females.

To determine the GRNs driving fibrogenic phenotypes, we compared enrichment of eRegulons across major cell populations of Saline and AngII mouse hearts (Figure 3E-F). Among the top eRegulons differentially enriched between the two conditions were those such as Smad3, a key mediator of TGFβ signaling, Egr3, Nfkb1, and Nfatc2, all implicated in fibrosis^13–15^; (Figure 3E). Also, among enriched eRegulons were those that are less recognized in cardiac fibrosis such as Creb3l2, and Etv6. eRegulons with the most marked reduction in enrichment included androgen receptor (Ar), Cebpd, *Krüppel-like factor* (KLF) family members (including Klf2, 4, 6 and 9)*, Jun* family members (Jun, Junb, Jund) and Tcf21. Many of these, including Cebpd and Klf4, have been independently demonstrated to be downregulated upon AngII stimulation of the heart and other models of fibrosis ^16,17^. Finally, evaluation of potential sex-specific GRNs showed the majority of eRegulons were present in both sexes (Figure 3G).

### *Elevated Cilp* and *Thbs4* expression define fibrogenic fibroblasts in mouse and human non-ischemic cardiac fibrosis

Analysis of AngII-treated mouse hearts revealed an increase in proportion of a fibroblast population enriched for fibrosis-related gene transcripts. As identified in our previous study ^10^, we found these cells highly enriched for *Cilp* and *Thbs4* transcripts (Figure 4). Here, we collectively refer to these cells as ‘Fibro-Cilp’, and as previously reported^10^, were able to determine their presence in regions of fibrosis within hearts of AngII-treated mice (Figure 4A-C and Supplementary Figure 7). These cells are present in both female and male contexts^10^. To evaluate *Cilp* and *Thsbs4* as markers of fibrogenic cells in a different model of non-ischemic fibrosis, we examined data from mice 18 days after thoracic aorta constriction (TAC)^18^. Similar to our AngII model, fibrogenic cells of this model were highly enriched for *Thbs4* and *Cilp* transcripts, with THBS4 detectable within fibrotic regions of TAC mouse hearts^19^ (Figure 4D-F and Supplementary Figure 8). We next determined whether Fibro-Cilp cells were present within the dataset prepared by multiomic analysis of the human heart (Figure 2). Here we noted fibroblasts were comprised of two cell subsets: one subset that is highly enriched for fibrogenic gene signatures and another that is not (Figure 4G-H). Corresponding to our analysis of AngII and TAC mouse datasets, we found that fibrogenic cells in our human dataset are enriched for CILP and THBS4 transcripts (Figure 4G-H and Supplementary Figures 9 and 10), with THBS4 detectable within the human heart tissue (Figure 4I and Supplementary Figure 10C). Consistent with these findings, we also found high enrichment of THBS4 transcript within remote fibrosis regions of human hearts following MI, determined using a dataset from an unaffiliated study^20^ (Figure 4J). Together our analysis reveals a common fibrogenic cell type that emerges in the context of non-ischemic fibrosis, irrespective of species or biological sex.

**Figure 4.**
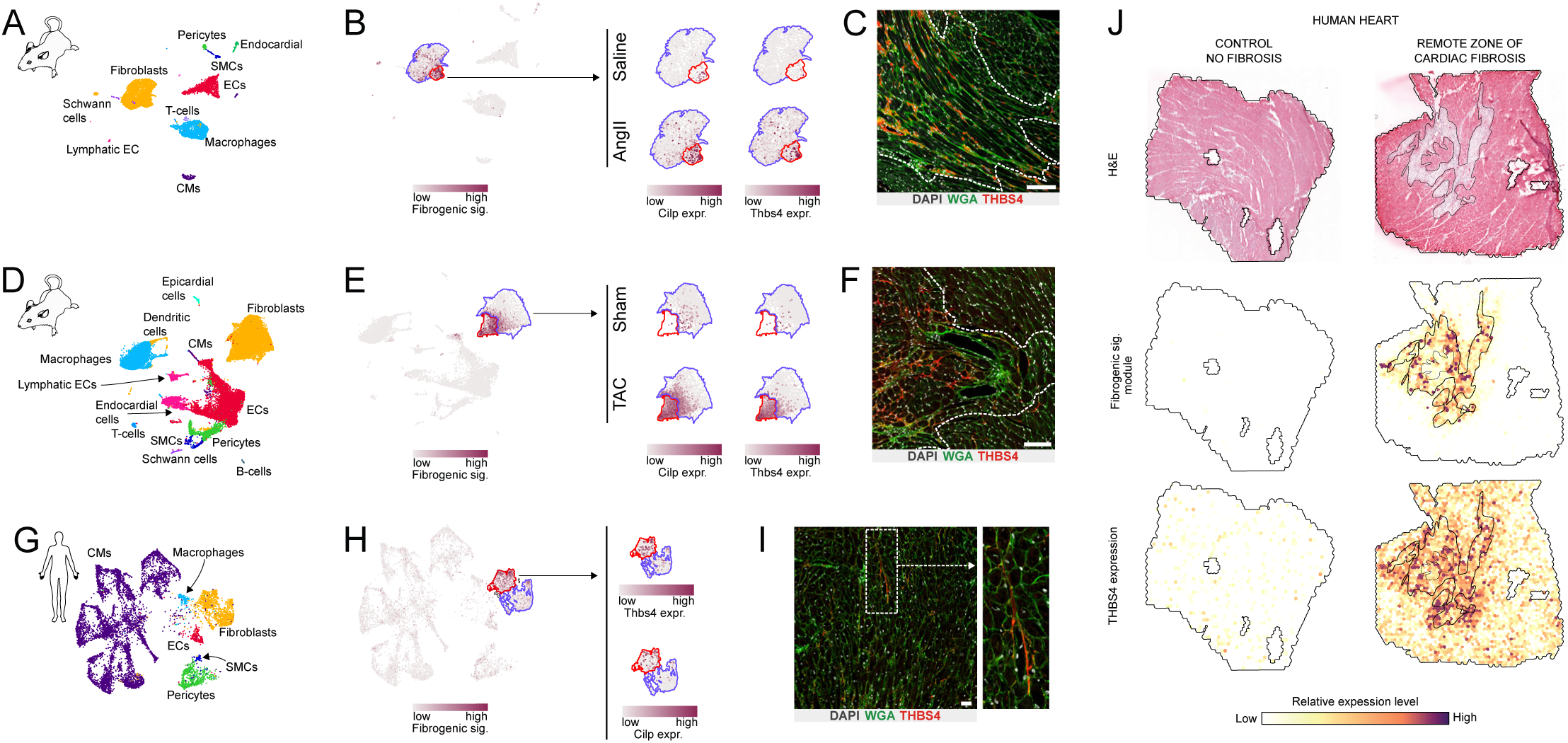
Fibro-Cilp cells are present in contexts of mouse and human non-ischemic cardiac fibrosis. A. UMAP of mouse cellulome developed using the multiomic dataset generated in this study. B. UMAP showing Fibro-Cilp subcluster of fibroblasts and corresponding enrichment of transcripts corresponding to a fibrogenic signature (formed by the sum of average transcripts/cell for Col1a1, Col3a1, Col8a1, Tnc and Postn genes). Right-most panels show fibroblasts UMAPs with heatmaps of Cilp and Thbs4 gene transcripts in context of Saline and AngII treatment. C. Immunostaining of AngII mouse hearts for THBS4. Nuclei and cell boundaries counterstained with DAPI and wheat germ agglutinin (WGA), respectively. Scale bar indicates 100 µm. D. As for A, but using TAC dataset (GSE155882 from ref. ^21^ ). E. As for B, but using TAC dataset (GSE155882 from ref. ^21^ ). F. Immunostaining of TAC mouse hearts for THBS4. Nuclei and cell boundaries counterstained with DAPI and wheat germ agglutinin (WGA), respectively. Scale bar indicates 100 µm. G. UMAP of human cellulome developed using the multiomic dataset generated in this study. H. UMAP of human cellulome showing Fibro-Cilp subcluster of fibroblasts and corresponding enrichment of transcripts corresponding to a fibrogenic signature (as defined above). Right-most panels indicate Thbs4 and Cilp gene transcript enrichment within the Fibro-Cilp subcluster. I. Immunostaining of donor human heart for THBS4. Nuclei and cell boundaries counterstained with DAPI and wheat germ agglutinin (WGA), respectively. Scale bar indicates 100 µm. J. Spatial transcriptomic mapping of remote regions of human cardiac cross sections highlighting the fibrogenic signature (as defined above) and *THBS4* expression in control samples (from individual not undergone myocardial infarction) and remote zones of cardiac fibrosis (from individuals who have had myocardial infarction; from ref. ^20^.

### Multi-model and multiomic single-cell analyses identify core TFs shaping Fibro-Cilp phenotype

Next, we focused on identifying the gene regulatory circuits driving Fibro-Cilp phenotype. To do this, we initially compared Fibro-Cilp to other fibroblasts from AngII mouse hearts. As noted previously (Figure 4), we found high enrichment of transcripts corresponding to genes linked to fibrosis within Fibro-Cilp cells. These included those forming a fibrosis signature (‘fibrosis signature’ genes: *Col1a1, Col3a1, Col8a1, Postn,* and *Tnc*) in addition to other genes including *Eln* and *Piezo2* (Figure 5A-B; Supplementary Table 3). Accordingly, top GO terms representing genes enriched in Fibro-Cilp included those associated with ECM (Figure 5C).

**Figure 5.**
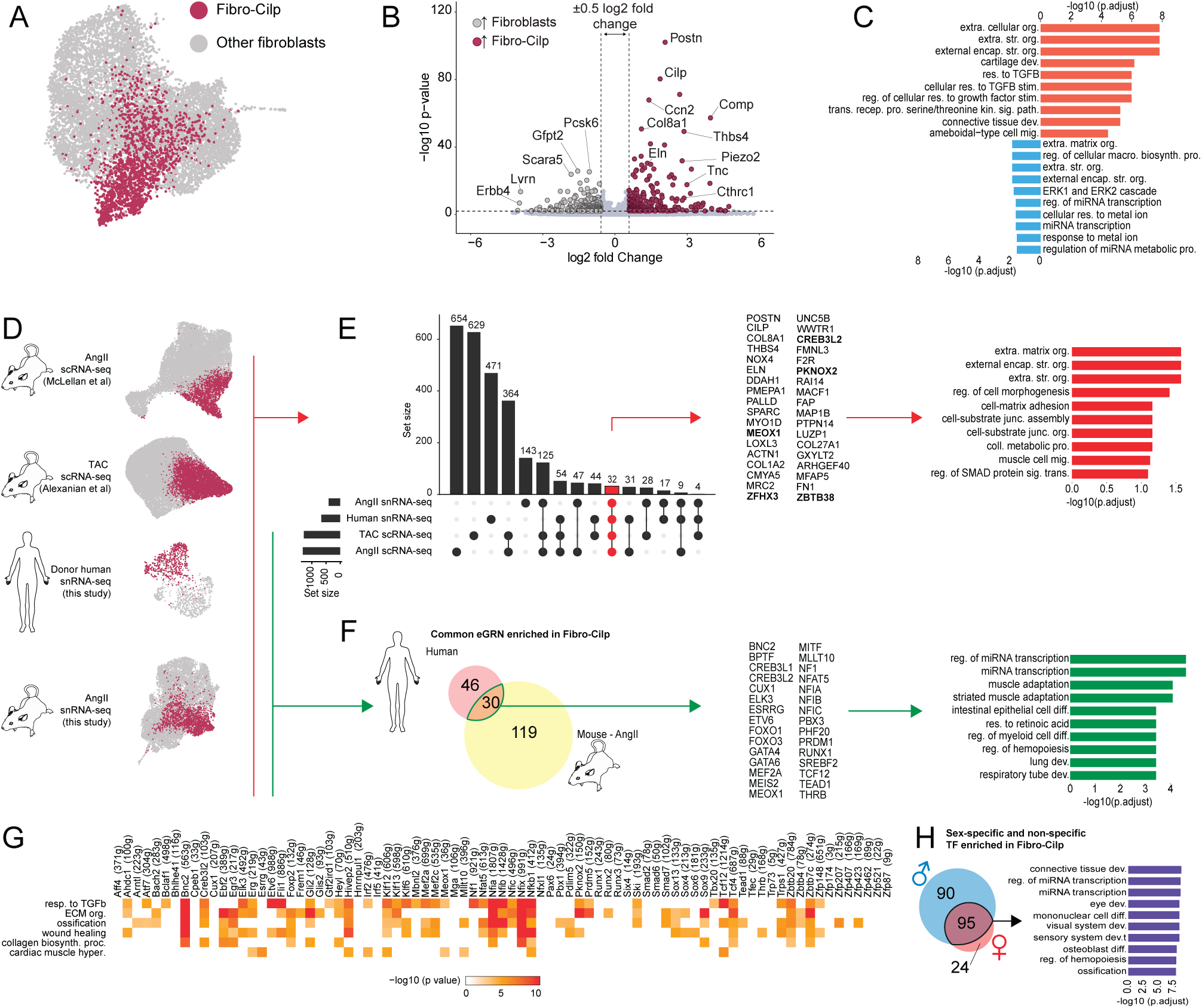
Common phenotype of Fibro-Cilp cells in mouse and human hearts are governed by shared gene regulatory circuits. A. UMAP projection of cardiac fibroblasts from Saline and AngII mouse hearts used for subsequent analyses in B and C. B. Volcano plot highlighting the up- and downregulated genes in Fibro-Cilp cells when compared to other fibroblasts from AngII mouse hearts. C. Top-10 GO terms corresponding to genes up- or downregulated in Fibro-Cilp cells compared to other fibroblasts (orange and blue bars, respectively). D. UMAPs of fibroblasts from mouse and human cardiac single-cell genomic datasets. Fibro-Cilp and other fibroblasts (non-Fibro-Cilp cells) shown in red and grey, respectively. E. Overlap of genes enriched in Fibro-Cilp cells compared to other fibroblasts from the datasets shown in D. Upset plot (left-most panel) summarized entire gene sets from each datasets (horizontal bars) and genes that are present uniquely or in multiple datasets (vertical bars). Genes present in all datasets (indicated in red; 32 genes) and listed (middle panel; bolded genes indicate TFs). Top 10 enriched GO terms corresponding to this list is shown in right-most panel. F. Overlap in eRegulons identified as enriched in Fibro-Cilp cells vs other fibroblasts from mouse AngII and human donor datasets. Overlapping TF list and top GO terms corresponding to the TF genes are shown in middle and far-right panels, respectively. G. Heat map of GO enrichment of TF target genes within eRegulons corresponding to fibrosis- and hypertrophy-related terms. H. Overlap of lists of eRegulons enriched in Fibro-Cilp vs other fibroblasts from AngII mice independently determined for each sex.

To assess the generalizability of the Fibro-Cilp phenotype, we drew on combined data from this study, our previous scRNA-seq study of AngII-induced fibrosis^10^, TAC mouse hearts^21^ (Figure 5D). Differential gene expression analysis (p<0.01) comparing Fibro-Cilp to other fibroblasts confirmed similar features of Fibro-Cilp cells from different non-ischemic contexts. There was high level of overlap between Fibro-Cilp-enriched gene sets from AngII and TAC-stressed heart scRNA-seq datasets (Figure 5E). There was markedly less overlap between Fibro-Cilp-enriched genes from scRNA-seq and snRNA-seq datasets from AngII-treated mouse hearts, reflecting the reduced number of genes captured by the snRNA-seq approach. However, we identified 32 genes—including fibrosis-linked matricellular genes *Postn, Cilp, Thbs4, Ddah2, Sparc* (Supplementary Table 4)—that are highly enriched in Fibro-Cilp cells in all four contexts. GO enrichment analysis showed these were primarily linked to ECM- and fibrosis-related programs (Figure 5E).

Next, we leveraged the human and mouse multiomic datasets to identify eRegulons that are highly enriched in Fibro-Cilp cells compared to other fibroblasts across both species. Our analysis found 30 common TFs more active in Fibro-Cilp, including many TFs that have been recently associated with fibrosis (Figure 5F; Supplementary Table 4). These included MEOX1^18^, TEAD1^9^, RUNX1^22,23^ amongst others. We also discovered many TFs that have not—to the best of our knowledge—been associated with fibrosis. GO enrichment analysis of these identified the disparate functions these TFs participate in, with many top statistically significant terms linked to tissue development or differentiation (Figure 5F)

We also found genes comprising eRegulons varied, indicative of the different aspects of the Fibro-Cilp phenotype they regulate. While the numbers of genes linked to a TF is a function of statistical parameters applied to undertake eRegulon discovery^12^, the largest eRegulons discovered by our analysis of AngII hearts were Nfia, Nfib, and Tcf12, associated with 1807, 1428 and 1214 genes, respectively (Figure 5G). To infer the potential role of TFs in establishing the Fibro-Cilp phenotype, we performed GO enrichment analysis on genes constituting eRegulons. Consistent with many genes upregulated in fibroblasts being linked to TGFβ responses (Figure 5C), 47% of TFs had statistically significant enrichment of genes linked to the *response to TGFβ* GO term. Further, many TF-linked genes were involved with terms linked to ECM remodeling, with some linked to cardiac hypertrophy (Figure 5G).

To study potential sex differences, we determined eRegulons that distinguish Fibro-Cilp from fibroblasts from AngII-treated female and male mice separately (Figure 5H). We found most Fibro-Cilp-enriched eRegulons in female mouse hearts were those enriched in Fibro-Cilp cells in male hearts, while we found more that were unique to males, based on our statistical parameters (Figure 5H). Nevertheless, we identified 95 TFs that are highly active in Fibro-Cilp cells in both sexes with these corresponding gene programs linked to ECM remodeling such as ‘*connective tissue development*’, ‘*osteoblast differentiation*’ and ‘*ossification*’. Together, our analyses have identified core TFs that drive Fibro-Cilp’s phenotype irrespective of species, biological sex or model of non-ischemic fibrosis.

### Reverse cardiac remodeling identifies transcription factors driving reversible non-ischemic fibrosis

Next, we sought to determine the plasticity and elasticity of GRNs driving cardiac fibrosis. To achieve this, we analyzed hearts of AngII and AngII+Rec mice (Figure 1A). We found withdrawal of AngII infusion led to reduction of fibrosis—determined by PSR staining of tissue and subsequent quantification (Figure 6A)— associated with cessation of AngII-induced hypertension (Supplementary Figure 11). Further examination of gene expression in fibroblasts of AngII+Rec compared to AngII mice showed downregulation of several fibrosis-associated genes including those encoding ECM components and key matricular protein genes such as *Postn*, *Cilp,* and *Thbs4* (Figure 6B). Consistent with a reduction Fibro-Cilp marker gene transcripts (*Cilp* and *Thbs4*), importantly, we found a reduction of this fibroblast subset within total cardiac fibroblasts. This was independent of sex, and resulted in AngII+Rec Fibro-Cilp cells reaching levels comparable to Saline mice (Figure 6C). Finally, reduction of Fibro-Cilp cells and *Thbs4* gene expression was validated by immunostaining which confirmed reduced fibrotic regions and THBS4 within hearts after recovery from AngII infusion (Figure 6D-E).

**Figure 6.**
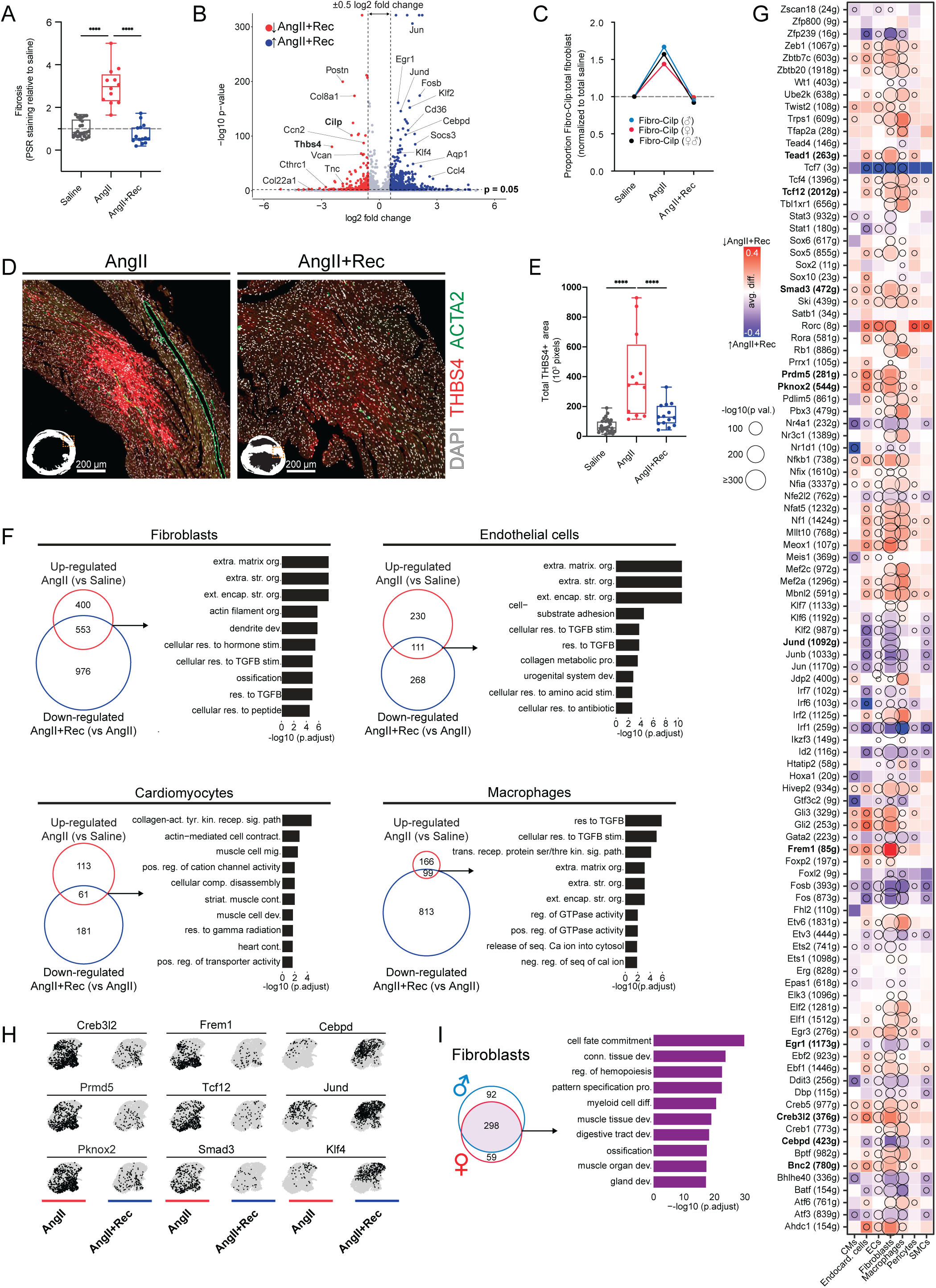
AngII withdrawal results in fibrosis regression, identifying TFs driving cardiac fibrosis. A. Cardiac fibrosis in AngII and AngII+Rec mouse hearts relative to Saline mouse hearts determined by quantifying picrosirius red staining. ****p < 0.0001 (one-way ANOVA). Dotted line indicates mean (Saline). B. Volcano plot of genes increased (blue) or decreased in gene expression in other fibroblasts from AngII+Rec mouse hearts compared to AngII mouse hearts. Note reduction in Thbs4 and Cilp gene expression. C. Cellular abundances of Fibro-Cilp cells relative to total cells in the dataset corresponding to Saline, AngII and AngII+Rec groups. Proportion data is normalized to Saline group. Data specific to females (red dot/lines), males (blue dots/lines) or both female and males (black dots/lines) are shown. D. Representative micrographs of THBS4 immunostaining within AngII and AngII+Rec mouse heart sections. E. Change in THBS4 immunostaining in cardiac cross sections from AngII and AngII+Rec mouse hearts (N= 5-6 animals per group). ****p < 0.0001 (one-way ANOVA test). F. Overlap of genes upregulated in major cardiac cells populations from AngII mouse hearts (relative to Saline) and those that are downregulated in AngII+Rec mouse hearts (relatively to AngII). Venn diagrams summarize unique and common genes within these gene sets. Bar plots summarize GO terms corresponding to overlapping genes. G. Heatmap/dot plot summarizing changes in eRegulon enrichment in cardiac cell types from AngII and AngII+Rec mouse hearts. H. UMAPs displaying binarized eRegulon enrichment within fibroblasts from AngII or AngII+Rec mouse heart datasets. First two columns (Creb3l2, Prmd5, Pknox2, Frem1, Tcf12 and Smad3) show examples of TFs that are more active in AngII mouse heart fibroblasts (i.e. they are repressed in AngII+Rec mouse heart fibroblasts). Third column (Cepbd, Jund, Klf4) show examples of TFs that are repressed in AngII mouse hearts and are robustly activated in AngII+Rec mouse heart fibroblasts. I. Summary of overlap eRegulons that are more highly enriched in cardiac fibroblasts in AngII+Rec compared to AngII mouse hearts, from each sex. Venn diagram summarizes sex-specific and non-specific eRegulons. Bar plot summarizes top-10 GO terms related to TFs comprising overlapping eRegulons.

To determine the extent to which the fibrosis-inducing gene signature is reversed, we compared genes upregulated in AngII (AngII vs Saline) and then downregulated in recovery (AngII+Rec vs AngII) in major cardiac cell types. GO enrichment analysis revealed multiple processes associated with ECM and fibrosis (Figure 6F) reversibly induced in fibroblasts, ECs and macrophages. Further, top reversibly induced genes in CMs correlated with processes involved in ECM signaling/interaction or those involved in myocyte structure (Figure 6F).

Next, we examined whether the reversal of fibrosis in AngII+Rec mice corresponds to a reversal of fibrosis-inducing TF activity. Comparison of eRegulon enrichment in AngII and AngII+Rec cell types revealed induction or repression of TF activity consistent with recovery from fibrosis. The activity of fibrogenic TFs induced by AngII such as Smad3—reflecting TGFβ signaling—Frem1, Creb3l2, Meox1, Pknox and others showed corresponding down regulation in AngII-Rec (Figure 6G-H, Supplementary Figure 13 and 14). In contrast, many TF activities suppressed by AngII-infusion were increased in AngII+Rec cardiac cells. These included Cebpd, Jund, and Klf4 (Figure 6G-H, Supplementary Figure 14). Other TFs departed from this pattern, for example Id2—associated with repression of TGFβ signaling and fibrosis^24^—was absent from our list of top TFs suppressed by AngII, but was induced in AngII+Rec cells.

Finally, we compared changes in TF activity in fibroblasts of each sex after removal of AngII. Major TFs largely overlapped between sexes, and these common TFs corresponded to GO terms associated with ECM such as *ossification* and *connective tissue development* (Figure 6I, Supplementary Figure 12). These observations suggest core TFs governing fibrosis and its reversal are independent of sex.

### Validation of distinct roles of TFs in regulating fibroblast phenotype and ECM deposition

To validate the role of core TFs identified in the context of TGFβ signaling, we tested the impact of TF gene silencing on the proteome landscape of primary human ventricular fibroblasts treated with TGFβ (Figure 7, Supplementary Figure 15). TGFβ treatment is a well-established model of fibroblasts-myofibroblast differentiation. Although myofibroblasts are absent in the setting of fibrosis we have examined here, we chose this system because fibroblasts exhibit a TGFβ response gene signature in non-ischemic fibrosis (Figure 3).

**Figure 7.**
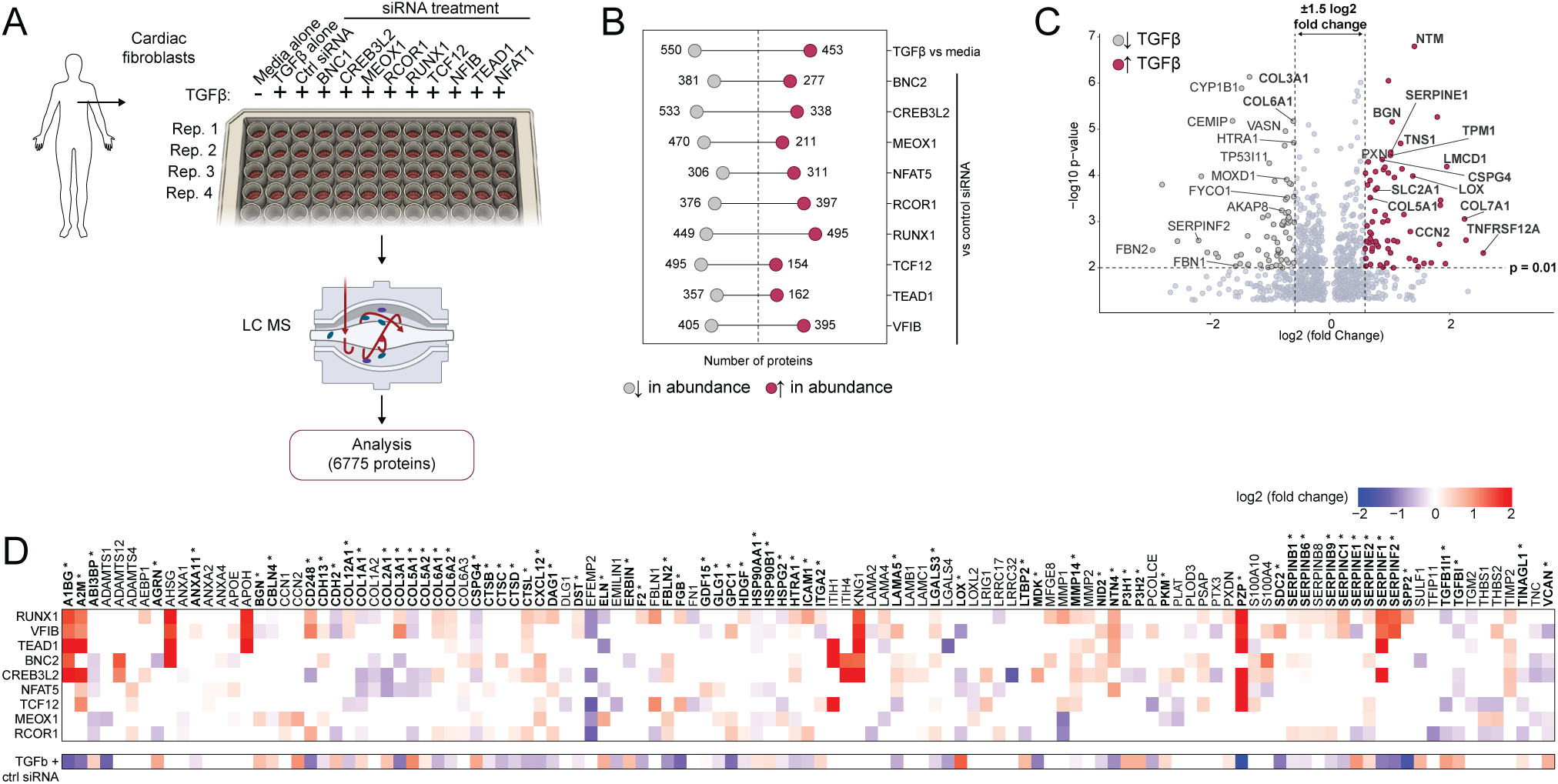
Proteomic validation of Fibro-Cilp-enriched TFs driving distinct facets of non-ischemic fibrosis. A. Experimental schematic for proteomic evaluation of roles of core TFs identified in this study in ECM production in TGFβ-treated human primary cardiac ventricular fibroblasts. All conditions were tested in four replicates. After TGFβ treatment, cell cultures were analyzed by high-throughput mass spectroscopy-based proteomic profiling before subsequent informatics analyses. B. Lollipop plots summarizing significant (p<0.05) differential expressed proteins in various comparisons indicated. TGFβ-treated cells were compared to media alone, whereas all TF siRNA-treated cells were compared to control siRNA-treated cells. C. Volcano plot summarizing proteins increased or decreased in abundance after TGFβ treatment, compared to media alone control. D. Heat map summarizing fold-increase or decrease in abundance of proteins in TF siRNA-treated cells compared to control siRNA-treated cells. Bottom panel shows protein abundance change in TGFβ + control (ctrl) siRNA vs media alone. Only ECM-related genes corresponding to GO term *Extracellular Matrix* (GO:0031012), with altered protein abundance levels in two or more conditions (control or TF siRNA groups) are displayed. Asterisks and bolded gene labels (x-axis) indicated proteins with TGFβ-induced protein abundance change being reversed by one or more siRNA targeting core TFs.

Treatment of cells with TGFβ, or TGFβ and siRNA corresponding to TFs, altered the proteomic landscape of fibroblasts (Figure 7B). Changes in several key protein expression patterns after TGFβ treatment were consistent with those anticipated based on transcriptional activity of fibroblasts in non-ischemic fibrosis. TGFβ-treatment elevated levels of CCN2, COL1A1 and COL5A2, and others, including those implicated in myofibroblast differentiation or fibrosis, while ECM-related proteins such as COL3A1—which are reduced in expression non-ischemic fibrosis—were also lower after TGFβ-treatment (Figure 7C). However, our proteomic profiling also found expression of hallmarks of non-ischemic fibrosis, such as COL8A1 and POSTN (Figure 7D), were reduced or unchanged following TGFβ treatment (Figure 7B, Supplementary Table 7). Therefore, *in vitro* assays, particularly those leading to myofibroblast differentiation, do not precisely recapitulate phenotypes observed in non-ischemic fibrosis *in vivo*.

Nevertheless, silencing of TFs revealed their distinct roles in regulating fibroblast phenotypes and ECM network (Figure 7C-D, Supplementary Table 7). For example, silencing CREB3L2, NFAT5, RCOR1 RUNX1 and NFIB all resulted in reduced levels of COL5A1 in response to TGFβ-treatment compared to unsilenced controls (Figure 7D). Distinct patterns of TF involvement were also found for other key ECM components such as COL1A1, COL3A1, COL5A2 and COL6A1. Similarly, TFs tested were important for expression of matricellular proteins, including CTHRC, TGFBI, TNC, and VCAN. CREB3L2 was a common, key regulator found to be important for expression of most of these genes. We also found evidence of potential gene repressive activities of TFs. For example, NFIB and RUNX1 silencing lead to a marked increase in COL3A1 protein levels, comparable to non-TGFβ treated controls (Figure 7D, Supplementary Table 7), while BNC2 and NFAT2 silencing had the opposite effect (Figure 7D). Together, these observations demonstrate the different roles that TFs play in fibroblast phenotypes and ECM deposition, and their complex roles in driving non-ischemic fibrosis.

## DISCUSSION

Fibrosis is a major contributor to cardiac dysfunction resulting in heart failure. Using mouse models, human heart tissue and leveraging single-cell transcriptomics and epigenomics, we aimed to gain a more comprehensive understanding of the GRNs driving cardiac cell identity and fibrogenic phenotypes. Our study has yielded new insights regarding the genetic circuits that are driving cell-specific and non-specific mechanisms of non-ischemic fibrosis.

A key feature of our study is the application of a paired multiomic approach—combining both snRNA-seq and snATAC-seq—to uncover transcriptional regulators of fibrosis development. While others have integrated these technologies using unpaired pipelines, where snRNA-seq and snATAC-seq libraries are prepared from separate pools of cells, in our approach transcripts and open chromatin fragments are extracted from the same nuclei for each cell. The paired approach overcomes challenges posed by integrating separate datasets, which relies on assumptions and data transformations such as converting snATAC-seq data to gene expression level. This may include the erroneous inference that a gene is active in a cell solely based on chromatin accessibility. These assumptions overly simplify the complex mechanisms underlying transcriptional regulation, and this is clearly demonstrated in our data. For example, snATAC-seq indicates the genetic locus of *Tcf21* is open in fibroblasts, SMCs and pericytes, despite *Tcf21* transcript being present only in fibroblasts. Using *Ppara* and *Sox9* as examples, our analyses also show that transcriptional activity (based on expression of target genes) does not always correlate with TF transcript level or chromatin accessibility of corresponding DNA-binding motifs. Armed with this unique dataset we calculated the major GRNs of single cells in different contexts and applied this to understand cellular diversity and drivers of fibrosis in different cardiac contexts.

A key application of single cell omics data is the classification of cell populations and subtypes based on distinct transcriptional signatures. GRNs identified in our analyses included TFs with well-established roles in cardiac cell development such as Nkx2-5, Fli1, Tcf21, Spi1, that are active in CMs, ECs, fibroblasts and macrophages, respectively, confirming the validity of our approach for the classification of major cardiac cell types. We were also able to recover TFs in a cell-specific manner from human data.

A major outcome of our analysis was the determination of key TFs that drive non-ischemic cardiac fibrosis. Most TFs exhibiting altered activity following AngII treatment operated in multiple cell-types. Shown here and previously^10^, virtually all cardiac cell types contributed to ECM remodeling with gene programs associated with TGFβ signaling. Reflecting this, Smad3—a key mediator of TGFβ signaling—was activated in multiple cell types. We also recovered a multitude of TFs that have been implicated in fibrosis including Nfkb1^15^, Bnc2^25^, Meox1^21^ and Egr3^13^. Further, we identified TFs with reduced activity upon development of fibrosis. These included JunD, Cebpd, Id2 and Klf4, all of which have been negatively correlated or implicated in dampening development of fibrosis^16,17,24,26,27^. Alongside TFs known to be involved in fibrosis our study uncovered several less-well associated, or previously unlinked, to cardiac fibrosis such as Creb3l2, Nfic and Phf20. Systematic exploration of these TFs will be required in the future to determine their role in the development of fibrosis.

Our analysis also provides a more comprehensive understanding of the highly fibrogenic cell population, Fibro-Cilp, that emerges in non-ischemic cardiac fibrosis. This population comprises both Fibroblast-*Cilp* and Fibroblast-*Thbs4* subsets we and others have recently identified^10,28^.

The generalized signature of Fibro-Cilp cells, independent of disease model, species or biological sex, is characterized by high expression of ECM structural genes and matricellular proteins. Although there were context dependent differences in these cells, GRN analysis identified a core set of TFs that regulated different aspects of Fibro-Cilp phenotypes. These included Meox1, Tead1, Runx1, and others which were recently identified as transcriptional regulators of ‘activated’ fibroblasts in single-cell genomic studies examining pressure-overloaded heart models^9,18^ or in the context of human cardiac remodeling^23^. Others we identified, such as Foxo1, have been demonstrated to regulate myofibroblast differentiation *in vitro* or *in vivo* following MI^29,30^. Using high-throughput proteomics, we were able to validate the roles of many of these in regulating human cardiac fibroblast ECM production *in vitro*.

We extended our exploration of key TFs driving fibrosis to cardiac recovery. Reverse remodeling of the heart has been noted in studies examining the impact of AngII or AngII receptor inhibitors, such as angiotensin II type 1 (AT_1_) blockers^31^, or murine pressure overload studies where bands constricting aortae or pulmonary arteries inducing cardiac hypertrophy are removed^32,33^. These studies demonstrate that both fibrosis and hypertrophy of the heart have the capacity for reversion to baseline levels. Here we confirm these findings showing that a key feature of reverse remodeling elicited by AngII withdrawal is the elimination of fibrosis, which occurs regardless of sex.

More recently, reverse remodeling has been explored using single-cell genomics. Alexanian and colleagues^21^ showed *bromodomain and extra-terminal domain* (BET) inhibitor JQ1 reverses pressure overload-induced cardiac remodeling. In this model *Meox1* is a key mediator of fibroblast activation that is downregulated upon JQ1 triggered recovery. Consistent with this, our analysis shows *Meox1* activation under AngII treatment and subsequent downregulation during recovery. Although we found that *Meox1* was not only important for fibroblast activation. *Meox1* was present in fibroblast populations enriched for *Wif1*^3,34,35^, in non-stressed hearts and a range of cell types—including macrophages, ECs, endocardial cells, and to a lesser degree, CMs—following AngII treatment. Another TF, *Runx1*, is implicated in reverse remodeling following LVAD implantation^36^. This too, we found, was reversibly activated in fibroblasts (and other cells), although it was not amongst the top significantly activated and deactivated TFs after AngII treatment and withdrawal (respectively) in our analysis. Among the top genes downregulated under AngII and increased in expression following withdrawal was Klf4, Cebpd and Id2, which are implicated in attenuating cardiac fibrosis^16,17,37^. KLF4 also controls other aspects of cell activity that may be related to controlling fibroblast phenotypes, including epigenetic programming^27^. Overall, our data helps tease-out the mechanisms and key TFs driving not just detrimental fibrosis but those driving recovery.

In this study we focused on transcriptional circuits driving fibrosis. We also restricted our consideration to activators of gene transcription rather than repressors. The^38^ approach we have applied here, and our dataset, could be expanded to these. Findings could also be extended to the examination of other cardiac phenotypes aside from fibrosis, such as CM hypertrophy, as has been examined by others^38^. Sex-specific patterns of gene expression and corresponding GRNs were also beyond the scope of this study and will explored in subsequent studies by us and others.

Finally, there are several limitations of our study which may have restricted our ability to fully uncover TFs driving cell identity and fibrogenic responses. These include: low yield of reads from our transcriptomic and epigenomic libraries leading to reduced power to discover eRegulons; low numbers of cells of specific cell types; and heterogeneity of sampling sites of human cardiac tissue. Further, in all our data, the reduced capture of transcripts due to nuclei isolation and permeabilization protocols, may have reduced our capacity to discover and identify eRegulons, which are partly contingent on capturing reads corresponding to TFs^12^.

In summary, by integrating data from multiple mouse models, human samples and leveraging single-cell multiomic methodologies, we have captured key TFs that drive cardiac fibrosis. In addition to identifying core transcriptional regulators that are well-implicated in cardiac fibrosis, we have also discovered a multitude of new TFs, involved both in induction and reversal of fibrosis. Collectively, this work provides exciting new avenues of research for informing clinical applications for reversing fibrosis and its deleterious effects.

## Methods

All experimental procedures were performed on C57BL/6J mice following protocols approved by Alfred Research Alliance (ARA) Animal Ethics Committee (application number E/1990/2020/B). Human studies were approved by the Alfred Hospital Ethics Committee (application number 143/19). The Sydney Heart Bank at the University of Sydney has Human Research Ethics Approval 122/2021.

For detailed additional information see Supplementary Material.

## Acknowledgements

The authors acknowledge use of the facilities at the Alfred Research Alliance Flowcore Cytometry Platform, the Monash Histology Platform, and the Baker Heart and Diabetes Institute Microscopy Platform. The authors acknowledge the Australian Red Cross for the historical coordination of the donor heart collections.

This work was funded by grants from the National Health and Medical Research Council (NHMRC; GNT1188503, GNT2021463 ) to A.R.P., and Australian Government Defense Science Institute RhD Student Grant to T.L.G.. Further, T.L.G., C.D.C., C.K., and G.E.F., were Supported by Australian Government Research Training Program stipends. A.R.P. is the recipient of the National Heart Foundation Australia, Future Leader Fellow, Level 3 (reference number 107335). D.W.G was supported by the National Heart Foundation Australia (reference number 105072) and by grants from the Medical Research Future Fund (MRFF; MRF1201805, MRF2024039). J.R.M was supported by a NHMRC Senior Research Fellowship (grant number 1078985) and Baker Fellowship (The Baker Foundation, Australia).

A provisional patent application has been filed based on the research findings described here.

## SUPPLEMENTARY MATERIAL

### Materials and Methods

#### Statement regarding human specimens

Human specimens involved in this study were obtained from the Sydney Heart Biobank of the University of Sydney under a Material Transfers Agreement (Ref: #CT25371) ^39^. This study complies with the regulations and ethical practices within the Alfred Hospital Ethics Committee and was approved under project number 143/19. In total 4 male and 4 female donor samples were obtained. The donor hearts were not used for heart transplantation because of logistical reasons at the time and are confirmed on formal anatomical pathology examination to be histologically normal.

#### Immunohistochemical analyses of human heart tissue

Human heart samples were embedded in OCT and sectioned onto SuperFrost slides at 10 µm thickness then stored at -80°C. For immunohistochemical analysis, slides were placed in -20°C for 10 minutes, then sections were fixed using cold 4% paraformaldehyde (PFA) in PBS at pH 6.9 for 10 minutes then permeabilized with 0.5% Triton-X (T9284, Sigma) in PBS for 10 minutes. Sections were then blocked with goat-block solution (2% goat serum [G9023, Sigma], 1% BSA [A3608, Sigma], 0.1% cold fish skin gelatin [G7765, Sigma], 0.1% Triton X-100). Primary antibodies were prepared in goat-block solution and incubated for 3 hours at r.t. followed by 3×5-minute washes with 0.05% Tween-20 in PBS (PBS-T). Incubation of secondary antibodies prepared in PBS-T was done at r.t. for 1 hour. Secondary antibodies were washed with 2×5-minute washes of 1 µg/mL WGA (29022-1, Biotium) in PBS-T followed by a 10-minute wash with 1 µg/mL DAPI in PBS-T. Sections were mounted and imaged using a Nikon A1r confocal microscope with a CFI Plan Apochromat VC 20× magnification lens of numerical aperture 0.75. Note, for anti-THBS4 staining, an additional incubation period step is required were the biotinylated antibody, prepared in PBS-T, is incubated for 1 hour at r.t. post primary incubation then washed 3× for 5-minutes with PBS-T. List of antibodies and dyes can be found in Supplementary Table 8 and 9.

#### Preparation of human heart samples for snRNA and snATAC-sequencing

Approximately 5 mg of human heart sample per subject was placed on dry ice from liquid nitrogen storage. As the tissue equilibrated in temperature on dry ice, a single stainless-steel bead (69989, QIAGEN) was placed in a safe-lock tube (0030120094, Eppendorf®), followed by the tissue, and 250 µL of cold *NP Buffer* (Supplementary Table 8 and 9) ^40^. Tissues were homogenized using the TissueLyser II (85300, QIAGEN) with four 10-second pulses at 25 Hz. At the start of the homogenization, a 15-minute timer was set. Post homogenization, an additional 250 µL of cold *NP Buffer* was added to each sample and samples were left on ice (4°C) till the end of the 15-minute incubation. Suspension was filtered through 20 µm cell strainers (43-50020-03, pluriSelect) into 50 mL LoBind tubes (0030122232 Eppendorf®) followed by 500 µL of cold *Fx Buffer* (Supplementary Table 8 and 9). Nuclei were pelleted at 1,000×*g*, for 5 minutes at 4°C. The supernatant was aspirated, and nuclei were gently resuspended in 100 µL cold *Fx Buffer*. For staining of cardiomyocyte nuclei, we conjugated anti-PCM1 rabbit antibody (Proteintech Group, 19856-1-AP) with fluorophore CF555 using and following manufacturer’s instructions of the Mix-n-Stain™ CF™ 555 Antibody Labeling Kit (MX555S20, Sigma-Aldrich). Nuclei suspensions were stained with anti-PCM1 and DAPI. Both DAPI+ (all non-myocyte nuclei) and PCM1+ DAPI+ nuclei (cardiomyocyte nuclei) populations were sorted separately using the BD FACS-ARIA I cell sorter at the ARA Flowcore. Nuclei were collected into dolphin microcentrifuge tubes (Z717533, Sorenson™) pre-coated with 5% BSA into 250 µL cold *Fx Buffer*.

#### Human transcriptomic and genomic library preparation

Post sorting, nuclei were pooled by equal proportions of nuclei from each subject (with 30% per sample contributing to cardiomyocyte nuclei). Nuclei permeabilization was performed as described by the 10xGenomics protocol *Nuclei Isolation from Complex Tissues for Single Cell Multiome ATAC + Gene Expression Sequencing* (10xGenomics, Doc# CG000375). Briefly, nuclei were spun down at 1,000×*g*, for 5 minutes at 4°C. The supernatant was carefully aspirated, and the pellet was resuspended in 100 µL of *0.1X Lysis Buffer* (10xGenomics, Doc# CG000375) for a 2-minute incubation on ice. After the incubation period, 1 mL of *Wash Buffer* (10xGenomics, Doc# CG000375) was added, and the nuclei were triturated 5 times. Finally, nuclei were pelleted at 1000×*g*, 5 minutes at 4°C and resuspended in *Diluted Nuclei Buffer* (10xGenomics, Doc# CG000375). A final nuclei count was performed, and the nuclei were filtered through the 40 µm Flowmi® cell strainers (H13680-0040, Bel-Art) prior to loading for partitioning. Transcriptomic (snRNA) and genomic (snATAC) libraries were prepared following *Chromium Next GEM Single Cell Multiome ATAC + Gene Expression Kits User Guide* with all required proprietary kits and equipment including the 10xGenomics Chromium Controller (1000204, 10xGenomics), Chromium Next GEM Single Cell Multiome ATAC + Gene Expression Kit (PN-1000283, 10xGenomics), Chromium Next GEM Chip J Single Cell Kit (PN-1000230, 10xGenomics) and Dual Index Kit TT Set A (PN-1000215, 10xGenomics).

#### Sequencing and pre-processing for human snRNA and snATAC libraries

Multiome ATAC FASTQ and GEX FASTQ files were downloaded from Illumina and were processed using Cell Ranger ARC v2.0.1 (10X Genomics) before downstream analysis. Reads were aligned to the pre-built human GRCh38 reference from 10X Genomics, arc-GRCh38-2020A-2.0.0, before ATAC and GEX matrices were constructed. Barcodes with high resolution in both matrices were then identified as a cell by joint cell calling algorithm. Summary for the Feature-barcode matrices were shown as follows.

|  | Sample a | Sample 1 | Sample 3 |
| --- | --- | --- | --- |
| Cells | 3,579 | 5,113 | 12,323 |
| Peaks | 159,234 | 112,293 | 126,495 |
| Gene | 27,421 | 29,826 | 32,173 |

Low quality cells in individual sample were filtered out based on the QC metrics using R v4.1 and v4.2, Signac v1.12.0 and Seurat v4.1.1/v5.0.1 (satijalab.org/seurat)^41,42^.

|  | Sample a | Sample 1 | Sample 3 |
| --- | --- | --- | --- |
| percent.mt | < 5 | <5 | <5 |
| nucleosome signal | <1.2 | <1.5 | <1.2 |
| TSS.enrichment | >2 | >1 | >2 |
| nCount_ATAC | < 200,000 & > 1,500 | < 150,000 & > 1,500 | < 150,000 & > 1,500 |
| nCount_RNA | < 250,000 & > 100 | < 15,000 & > 100 | < 20,000 & > 100 |
| Blacklist_ratio | <0.03 | <0.03 | <0.03 |

SCTransform^43^ was used to normalize the RNA data and regress out ribosomal protein genes (genes starting with Rps and Rpl) and mitochondrial genes to reduce technical artifacts. ATAC-seq data was normalized by term frequency inverse document frequency (TF-IDF). Three Seurat objects were created for each sample and cells were initially clustered using weighted nearest neighbor (WNN) method. Low quality clusters were removed based on nFeature_RNA, nCount_RNA and percent.mt. Remaining cells were re-clustered, and doublets were removed using DoubletFinder^44^ prior to sample integration.

#### DoubletFinder (Human)

|  | Sample a | Sample 1 | Sample 3 |
| --- | --- | --- | --- |
| Singlet | 2,830 | 3,175 | 7,819 |
| Doublet | 64 | 72 | 497 |

Seurat integration approach was performed with 3000 integration features used and canonical correlation analysis (CCA) was utilized to find the integration anchors. The anchors were selected based on the RNA assay and were used to correct the cell embedding for both RNA and ATAC data. Clustering was performed on integrated Seurat object using WNN method. Clusters lacking biological information were removed and remaining cells were re-clustered. Final dimensionality reduction was conducted using 40 principal components for RNA data, 40 corrected latent semantic indexing (LSI) for ATAC data and a resolution of 0.8 to explain variability not expected by chance to cluster the nuclei into cell populations. Uniform Maximum Approximation and Projection (UMAP) was used to visualize the data in two dimensions and ‘FindAllMarkers’ was used to identify cell clusters. The final dataset included 8,474 nuclei, 36,601 genes and 167,683 ATAC fragments across 3 samples.

#### Statement regarding animal experiments and husbandry

All animals and respective animal-related experiments were listed under ethics number E/1990/2020/B and approved by the Alfred Research Alliance (ARA) Animal Ethics Committee (AEC) in accordance with the National Health and Medical Research Council (NHMRC) of Australia. Animals were sourced from the Animal Resource Centre (ARC) located in Western Australia. Animals were maintained in 12-hour light/dark cycler with *ad libitum* access to food and water.

#### Experimental design

Hypertension-induced cardiac hypertrophy and fibrosis experiments were achieved by subcutaneous implantation of osmotic mini-pumps (Alzet, model 2002) loaded with 1.4 mg/kg/day angiotensin II (AngII) or saline (vehicle) to animals 10-12 weeks of age. All surgical procedures were conducted at Baker Heart and Diabetes institute by the Preclinical Cardiology Microsurgery and Imaging Platform (PCMIP). Alzet osmotic mini-pumps were removed after 2-weeks of AngII or saline in recovery cohort. All animals where Alzet osmotic mini-pump implants and explants were anesthetized using 1.7% isoflurane, with heart rates of 407 ± 97 beats/min. At experimental endpoint mice were anaesthetized with isoflurane with body temperature maintained at 37°C, and a 1.4F micro tipped transducer catheter (Millar, Houston TX) was used to measure arterial blood pressure (BP). Pressure catheterization procedures were performed by PCMIP at the Baker Heart and Diabetes Institute. Mice were euthanized by cardiac puncture and followed by perfusion outlined below for tissue collection.

#### Immunohistochemical analysis of mouse heart tissue

Animals were perfused with cold 200 mM KCl in PBS through the left ventricle until blood cleared, then fixed with 200 mM KCl ,4% PFA in PBS at pH = 6.9 for 10 minutes at 1 mL/min/mouse. Hearts were collected, fixed in 4% PFA in PBS at pH = 6.9 for 2 days at 4°C, and stored in 0.09% NaN_3_ in PBS at 4°C. With the assistance of the Monash Histology Platform (Monash University, Australia), hearts were processed, embedded in paraffin and sectioned onto SuperFrost slides at 4 µm thickness. Sections were subject to picrosirius red staining (PSR) and immunohistochemical analysis. For the detection of PPARα, GATA4, SOX9, SMA and THBS4, FFPE slides were heated to melt paraffin at 80°C for 1 minute, then incubated with 3 × 10-minute in xylene. Sections were rehydrated with 5-minute serial incubations of ethanol at 100%, 100%, 90% 70%, and 50%. Slides were introduced to water for 1 minute then rinsed with antigen retrieval buffer. Antigen retrieval was performed by placing the slides in 1 L of 1 × Tris-EDTA buffer (ab93684, Abcam) and microwaving at 1000W for 15 minutes. Once slides cooled, sections were permeabilized with 0.5% Triton-X and blocked for 1 hour at r.t. with goat block solution. Primary antibodies were prepared in goat block solution and incubated overnight at 4°C. Sections were washed with 3×5-minute washes with PBS-T then incubated with corresponding secondary antibody, prepared in PBS-T, for a minimum of 2 hours at r.t.. Secondary antibodies were washed with 2×5-minute washes of 1 µg/mL WGA in PBS-T followed by a 10-minute wash with 1 µg/mL DAPI in PBS-T. Sections were mounted and imaged using a Nikon A1r confocal microscope with a CFI Plan Apochromat VC 20× magnification lens of numerical aperture 0.75. Further PSR stained slides were scanned on the Aperio Scanscope AT Turbo while the Olympus VS200 Automated Slide Scanner was used for immunofluorescence slide scanning. Note, for anti-THBS4 staining, an additional incubation period step is required where the biotinylated antibody, prepared in PBS-T, is incubated for 2 hour at r.t. post primary incubation then washed 3× for 5-minutes with PBS-T. List of antibodies and dyes used is found in Supplementary Table 8 and 9.

#### Preparation of mouse heart tissue for snRNA and snATAC sequencing

Animals were injected with 100 U of heparin (1 U/µL prepared in saline) ∼30 minutes prior to lethabarb (300-400 mg/kg) injection. A modified *Langendorff* approach was taken, as previously described in McLellan *et al*. 2020^10^ and Farrugia *et al*. 2021^45^ to isolate viable cardiac myocytes (CMs) and cardiac non-myocytes. Briefly, hearts were perfused with cold *EDTA Buffer* then, using a combination of collagenases II and IV, and protease XIV, prepared in *Perfusion Buffer*, the hearts were dissociated through the LV with minimal mechanical aid to preserve the structure of the cardiomyocytes. Digestion was halted using *Stop Buffer* (Supplementary Table 8 and 9)^45^.

Non-myocytes were separated from CMs by sedimentation and gentle centrifugation at 100×*g* for 1 minute at 4°C. The non-myocytes, present in the supernatant, were separated and labelled with antibodies corresponding to CD45 and CD31, and stained with viability dyes SYTOX™ Green (SG) and Vybrant™ DyeCycle™ (VDR). Non-myocytes were sorted on the BD FACS-ARIA Fusion cell sorter at the ARA Flowcore. Single, viable (SG-VDR+) cells were sorted based on the presence/absence of CD45 and CD31 antigens; CD45+ (leukocytes; leuks), CD31+CD45-(endothelial cells; ECs), and CD45-CD3-(resident mesenchymal cells; RMCs). Non-myocytes were collected in dolphin microcentrifuge tubes pre-coated with 5% BSA and into 250 µL cold *Fx Buffer*.

For nuclei isolation of CMs (after separation from non-myocytes), CMs were washed with 5 mL of cold *Stop Buffer* and sedimented at 100×*g* for 1 minute at 4°C. The supernatant was discarded and the CMs were resuspended in 1 mL of cold *Stop Buffer* and transferred into a 5 mL tube snap-lock. They were spun one more time at the same conditions before resuspending the CM pellet in 750 µL of 2*× Hypo-osmotic solution* (Supplementary Table 8 and 9). The cells were triturated gently, then placed on ice for 5 minutes. Post incubation, samples were slowly triturated 3 times using a 5 mL Luer-lock syringe and a 30G needle. Samples were then pulled through 20 µM filters into 50 mL LoBind tubes followed by 500 µL of *Fx Buffer*. Nuclei were pelleted at 1,000×*g*, for 5 minutes at 4°C. CM nuclei were then stained with DAPI and subject to sorting on the BD FACS-ARIA I cell sorter at the ARA Flowcore. Nuclei were collected into dolphin microcentrifuge tubes pre-coated with 5% BSA into 250 µL cold *Fx Buffer*.

#### Mouse transcriptomic and genomic library preparation

Post sorting, non-myocyte cells from each group were pooled together at a representation of ∼35% ECs and 65% non-ECs (leuks and RMCs) per mouse, with 4 male and 4 female mice per group. Nuclei isolation was performed as described by the 10xGenomics protocol *Nuclei Isolation for Single Cell Multiome ATAC + Gene Expression Sequencing* (10xGenomics, Doc# CG000365). Briefly, cells were spun down at 400×*g*, for 5 minutes at 4°C. The supernatant was carefully aspirated, and the pellet was resuspended in 100 µL of chilled *Lysis Buffer* (10xGenomics, Doc# CG000365) for a 2-minute incubation on ice. After the incubation period, 1 mL of *Wash Buffer* (10xGenomics, Doc# CG000365) was added, and the nuclei were triturated 10 times. Nuclei were pelleted at 1000×*g*, 5 minutes at 4°C and resuspended in *Wash Buffer*. Two additional washes were performed, and non-CM nuclei were counted. At this step, CM nuclei were counted, and the CM and non-CM nuclei were combined to prepare one single nuclei suspension per experimental group containing 20% CM and 80% non-CM cardiac nuclei. Nuclei were pelleted and permeabilized following the *Nuclei Permeabilization* protocol as described (10xGenomics, Doc# CG000375). A final nuclei count was performed, and the nuclei were filtered through the 40 µm Flowmi® cell strainers prior to loading for partitioning. Transcriptomic (snRNA) and genomic (snATAC) libraries were prepared by loading 10,000 nuclei per lane and following *Chromium Next GEM Single Cell Multiome ATAC + Gene Expression Kits User Guide* with all required proprietary kits and equipment including the 10xGenomics Chromium Controller (1000204, 10xGenomics), Chromium Next GEM Single Cell Multiome ATAC + Gene Expression Kit (PN-1000283, 10xGenomics), Chromium Next GEM Chip J Single Cell Kit (PN-1000230, 10xGenomics) and Dual Index Kit TT Set A (PN-1000215, 10xGenomics).

#### Sequencing, pre-processing, QC, filtering and processing for mouse snRNA and snATAC libraries

Multiome ATAC FASTQ and GEX FASTQ files were downloaded from Illumina and was processed using Cell Ranger ARC v2.0.1 (10X Genomics) before downstream analysis. Reads were aligned to the pre-built mouse mm10 reference from 10X Genomics, arc-mm10-2020A-2.0.0, before ATAC and GEX matrices were constructed. Barcodes with high resolution in both matrices were then identified as a cell by joint cell calling algorithm. Summary for the Feature-barcode matrices were shown as below.

|  | Sample 1 (Saline) | Sample 2 (AngII) | Sample 3 (Recovery) |
| --- | --- | --- | --- |
| Cells | 8,283 | 9,439 | 13,265 |
| Peaks | 192,475 | 212,698 | 163,814 |
| Gene | 23,848 | 24,235 | 24,688 |

Low quality cells in individual sample were filtered out based on the QC metrics using R v4.1 and v4.2, Signac v1.12.0 and Seurat v4.1.1/v5.0.1 (satijalab.org/seurat).

|  | Sample 1 (Saline) | Sample 2 (AngII) | Sample 3 (Recovery) |
| --- | --- | --- | --- |
| percent.mt | < 40 | <50 | <30 |
| nucleosome_signal | <1.6 | <1.5 | <1.5 |
| TSS.enrichment | >1 | >1 | >1 |
| nCount_ATAC | < 125,000 & > 1,000 | < 150,000 & > 1,000 | < 125,000 & > 1,000 |
| nCount_RNA | < 9,000 & > 100 | < 8,500 & > 100 | < 7,500 & > 100 |

SCTransform^43^ was used to normalize the RNA data and regress out ribosomal protein genes (genes starting with Rps and Rpl) and mitochondrial genes to reduce technical artifacts. ATAC-seq data was normalized by term frequency inverse document frequency (TF-IDF). Three Seurat objects were created for each sample and cells were initially clustered using weighted nearest neighbor (WNN) method. Low quality clusters were removed based on nFeature_RNA, nCount_RNA and percent.mt.

Remaining cells were re-clustered, and doublets were removed using DoubletFinder^44^ prior to sample integration.

#### DoubletFinder

|  | Sample 1 (Saline) | Sample 2 (AngII) | Sample 3 (Recovery) |
| --- | --- | --- | --- |
| Singlet | 6,172 | 6,717 | 9,293 |
| Doublet | 264 | 383 | 383 |

Seurat integration approach was performed with 3000 integration features used and canonical correlation analysis (CCA) was utilized to find the integration anchors. The anchors were selected based on the RNA assay and were used to correct the cell embedding for both RNA and ATAC data. Clustering was performed on integrated Seurat object using WNN method. Dimensionality reduction was conducted using 40 principal components for RNA data, 40 corrected latent semantic indexing (LSI) for ATAC data and a resolution of 0.8 to explain variability not expected by chance to cluster the nuclei into cell populations. Uniform Maximum Approximation and Projection (UMAP) was used to visualize the data in two dimensions and ‘FindAllMarkers’ was used to identify cell clusters. The final dataset included 22,182 nuclei, 32,285 genes and 229,746 ATAC fragments across 3 samples comprising 6,717 nuclei from 2-week AngII, 6,172 nuclei from 2-week Saline and 9,293 from 2 weeks recovery.

#### Gene ontology analysis

GO enrichment analysis was conducted using the ‘enrichGO’ function within the ‘clusterProfiler’ R package version 4.4.4. Reference genomes obtained from http://geneontology.org were employed for all presented GO analyses. Enrichment analysis specifically for GO Biological Process terms (GO-BP) involved mapping our set of differentially expressed genes (adjusted *P* value < 0.01) to the *Mus musculus* background gene list. Statistically significant GO-BP terms were identified using a Benjamini-Hochberg adjusted *P value* cutoff of 0.01.

#### Incorporation of single-cell RNA sequencing data from publicly available data

To validate our findings from both the snRNA-seq and snATAC-seq datasets, we integrated differentially expressed genes obtained from publicly available scRNA-seq dataset (Reference: Alexanian et al.). Cells from this dataset were reclustered to isolated *Fibroblasts* and *Fibroblasts-Cilp/Thbs4* cells. Subsequently, a differential gene expression analysis was conducted between these two cell types. Genes with non-zero expression in >10% of cells in at least one of the experimental groups were included in the analysis. The ‘MASTcpmDetRate’ method, a modification of ‘MAST’, was employed for differential expression (DE) gene identification. A significance threshold of uncorrected *P* value < 0.01 was set unless otherwise specified. Upon completion of DE analysis for both the mouse and human snRNA-seq datasets as well as the publicly available dataset, a comparison of DE genes was performed using R packages, including RVenn (v1.1.0), VennDiagram (v1.7.3), and Venneuler (v1.1-3). Differentially expressed genes from all datasets were filtered based on P value threshold (P<0.01). Common differentially expressed genes were determined using the ‘overlap’ function within RVenn. Subsequently, Gene Ontology (GO) analysis was conducted on each overlapping gene list derived from this analysis, elucidating biologically relevant pathways specific to each set of overlapping genes.

#### Analysis of biologicals sex

To investigate the sexual dimorphic patters at transcriptional and epigenetic levels, male and female cells were isolated using scRNA-seq data based on the expression of Xist and five Y chromosome genes (Ddx3y, Eif2s3y, Gm29650, Kdm5d, and Uty), as outlined in Skelly et al. 2018^46^. In the initial step, a cell with non-zero expression of Xist but zero expression of our Y chromosome genes was categorized as a female cell. Conversely, a cell with non-zero expression for one or more of the five Y chromosome genes and zero expression of Xist was classified as a male cell. Moreover, to account for the fact that female cells should not contain a Y chromosome, and recognizing that male cells could express Xist transcripts, cells with low or moderate Xist expression without low expression of Y genes were further classified as male cells. Specifically, cells with summed Y expression greater than the lowest 10% of Y gene expression among male cells (as identified in step 1) and Xist expression less than the median Xist expression among female cells (as identified in step 1) were classified as male. Cells that expressed neither Xist nor Y chromosome genes or did not meet the specified criteria were excluded from the sex analysis. This analysis was conducted on pre-processed scRNA-seq data (quality controlled, normalized, and doublets removed) on per-sample basis, and the labels were transferred to the final Seurat object. As a result, approximately 60% of cells from our final dataset were assigned a sex for downstream sex analysis (6,708 female and 6,572 male cells).

#### Spatial Transcriptomic Data Analysis

Spatial transcriptomic analysis was carried out using published data from Kruppe *et al,* 2022^20^. Quality controlled spatial transcriptomic data was obtained from via cellxgene https://cellxgene.cziscience.com/collections/8191c283-0816-424b-9b61-c3e1d6258a77. Briefly, quality control processing involved exclusion of spots with less than 300 measured genes and less than 500 UMI, as well as filtering ribosomal and mitochondrial genes using Seurat (v4.0.1)^20^. sctTransform was used to normalize individual count matrices, with additional log-normalized (size-factor = 10,000) and scaled matrices calculated for comparative analysis. To assess our Fibro-Cilp signature and expression pattern of the regulon gene panels (see Table X), we selected the remote regions (RZ) from this myocardial infarction model, and further filtered sections based on fibrotic regions as seen with H&E staining. Module analysis was carried out for Fibro- (POSTN, COL1A1, COL3A1, COL8A1, TNC), Fibro-Cilp (THBS4, CILP, CTHRC1, SPARC), and all regulon gene lists using the Modules package (v0.12.0) (https://github.com/wahani/modules). Data is presented as a spatial feature plot, with outlined fibrotic region in the RZ tissue section.

#### Regulatory Analysis

##### SCENIC

To explore the dataset specific GRN, we firstly performed SCENIC v0.12, a package used to reconstruct the GRN from snRNA-seq^47^, on the RNA raw count matrix extracted from the final integrated Seurat object. Only genes with sum of expression over zero were remained for building GRN inference. A python script, arboreto_with_multiprocessing.py, from pyscenic package was used to generate a list of adjacencies between a potential transcription factor and its target genes with default GENIE3 algorithm. Next, regulons were predicted via *pyscenic ctx* from command line interface (CLI) with *--mask_dropouts* and *--nes_threshold* = 2 flags. GRNs were then built and scored by *pyscenic auc* from CLI with same NES threshold used for regulon prediction. Finally, 598 and 498 regulons were detected and imported into mouse and human Seurat objects respectively.

##### SCENIC+

To explore enhancer-driven GRNs (eGRNs), SCENIC+ version 1.0.1, a python package that builds eGRNs using combined snRNA-seq and snATAC-seq data^48^, was performed on the integrated dataset to identify the full dataset specific eRegulons. Pycistopic v1.0.3 was firstly used to initialize the cisTopic object and run the topic modelling using ATAC count matrix extracted from integrated Seurat object. Model with 38 regulatory topics was selected and used to infer candidate enhancer regions. Top 3000 regions per topic were used to binarize the topics. Once candidate enhancer regions were identified, pycistarget was used to find enriched motifs in these regions with customized parameter settings. We reduced the threshold used to calculate the AUC to 0.002, normalized enrichment score (NES) to 1.3, log2 fold change between the regions set and background to 0.5, and the minimum cis-regulatory model value to 2.

SCENIC+ object was then created using pre-processed snRNA data from integrated Seurat object, cisTopic object, and motif enrichment output from pycistarget. Cistromes was generated with standard procedure of SCENIC+ pipeline before building eGRNs. We used 150k upstream and downstream of the gene for the search space around the gene prior modelling enhancer-to-gene (R2G) correlation. To train the R2G model, random forest algorithm was used with training parameter n_estimators = 10000. For transcription factor to gene relationship (TF2G), a precomputed list of adjacencies that was generated using arboreto_with_multiprocessing.py from SCENIC ^47^ was used. Finally, eGRNs was built using build_grn() function with correlation coefficient threshold 0.02, GSEA permutation number 10000, and minimum number of target genes 3 used. In terms of human data, topic model with 38 topics was used and final SCENIC+ object was generated using a wrapper function run_scenicplus(). To identify saline and fibrogenic fibroblasts specific eGRNs, dataset was subset prior training R2G model and same approaches as mentioned before was used. However, we used correlation coefficient threshold 0.04 during eGRN building to capture more biological plausible eRegulons.

#### Fibroblast proteome reprogramming

Human primary cardiac fibroblasts (ventricle CC-2904, batch 20TL356511, single donor: female 37 years old) were cultured as described^49^. siRNA treatments were performed per well, 6 pmol siRNA in 25ml DMEM combined with 25ml DMEM containing 0.5ml RNA iMAX for 15 minutes, added to cells. After 24 hours of transfection, media was removed and replaced with DMEM + 0.5% FCS containing 10ng/mL TGFβ or saline control. 48 hours following TGFβ treatment, cells were washed with PBS and snap frozen. Proteomic sample preparation was performed using 1% SDS (pH 8), with protease and phosphatase inhibitor (Halt, Life Technologies, #78442) and proteomic preparation performed using the Sera-Mag-based workflow^50^. Spectra were acquired in data independent acquisition on an Q Exactive HF-X benchtop Orbitrap mass spectrometer coupled to an UltiMate™ NCS-3500RS nano-HPLC (Thermo Fisher Scientific) as described^51^. MS-based cellular proteomics data is deposited to the ProteomeXchange Consortium via the MASSive partner repository and available via MASSive with identifier (MSV000094580). For visualizing impact of TGFβ or siRNA on ECM-related genes, ECM-related gene list was created using Gene Ontology term 0031012 (*Extracellular Matrix*), and this list was further filtered for genes that were within the proteomic dataset. Proteins with altered abundance levels (relative to corresponding controls; p<0.05) in two or more conditions were presented as a heatmap.

### Supplementary Figures Legends

**Supplementary Figure 1.**
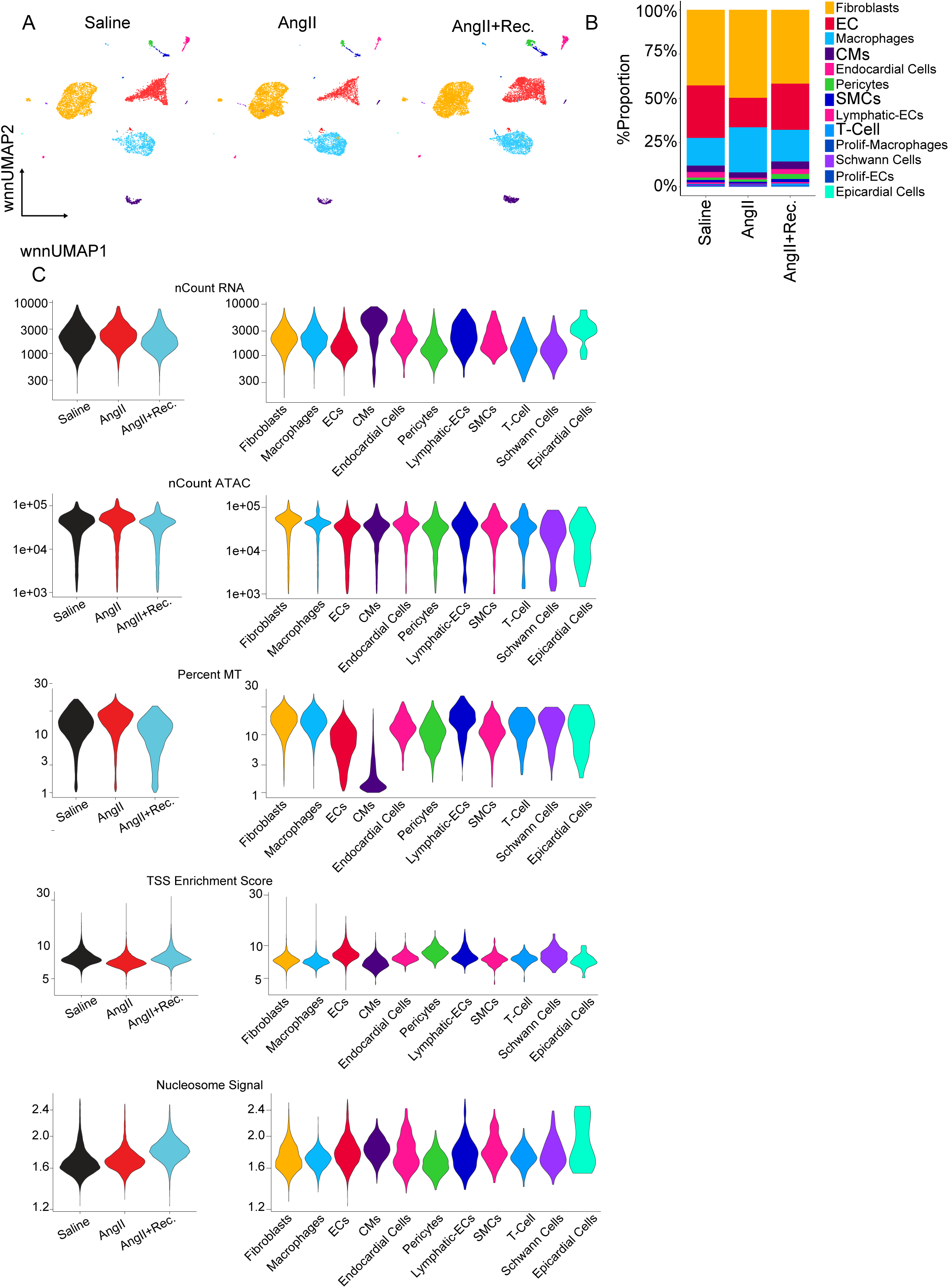
Sample and data quality of mouse paired-multiomic dataset (related to Figure 1). A. Weighted Nearest Neighbor (WNN) UMAP of samples from the three experimental conditions. B. Stacked bar plot summarizing relative proportions of cell types from different experimental conditions. C. Data quality parameters for samples from each condition or major cardiac cell types.

**Supplementary Figure 2.**
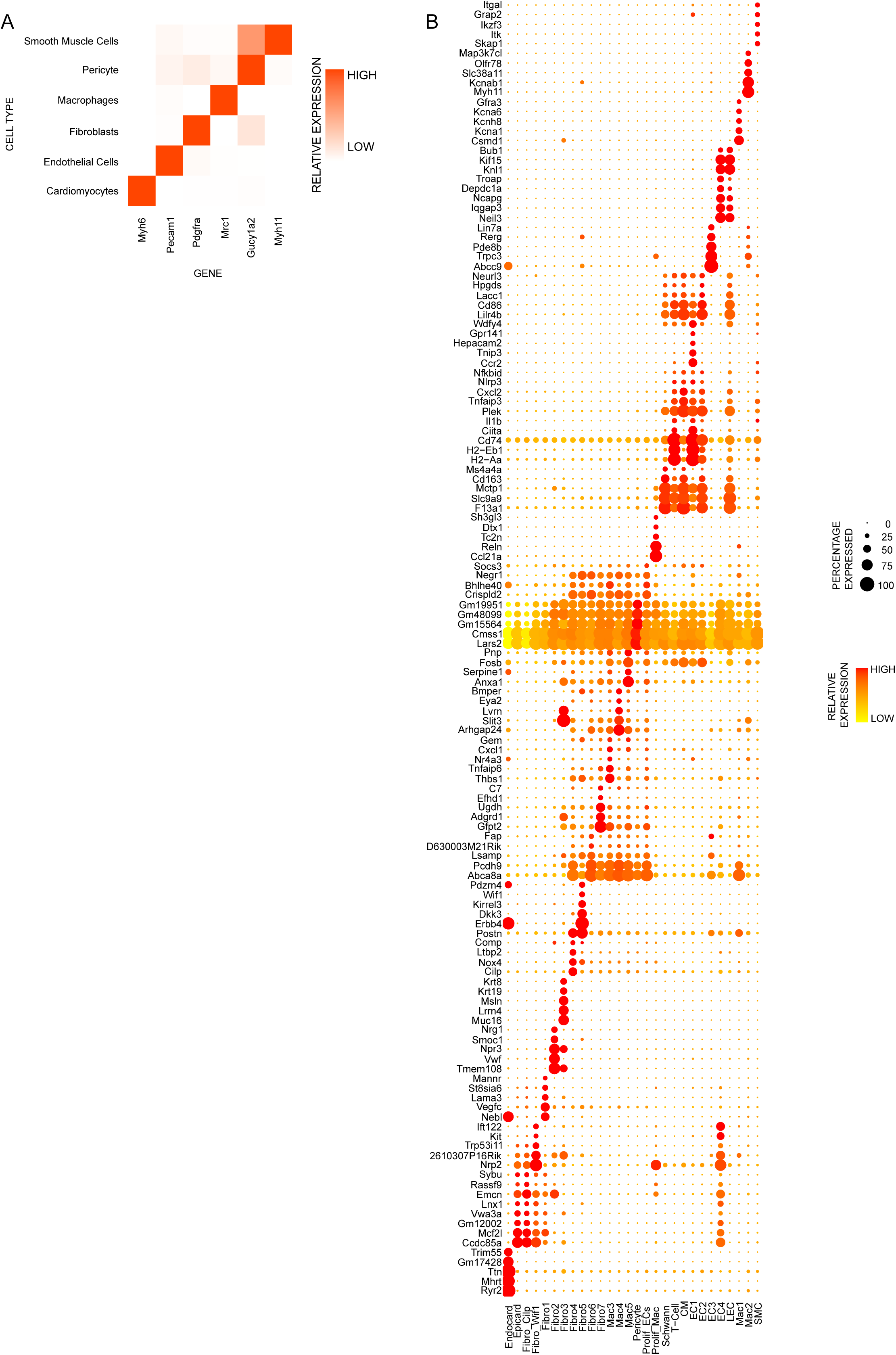
Markers of cardiac cell types within the mouse paired-multiomic dataset. A. Heat map of relative expression of canonical cell marker genes of major mouse cardiac cell types. B. Dot plot of the relative expression of the top 5 marker genes per cell sub-cluster. Dot size and color intensity indicate the proportion of cells expressing the genes relative average expression level, respectively, within each cell population.

**Supplementary Figure. 3.**
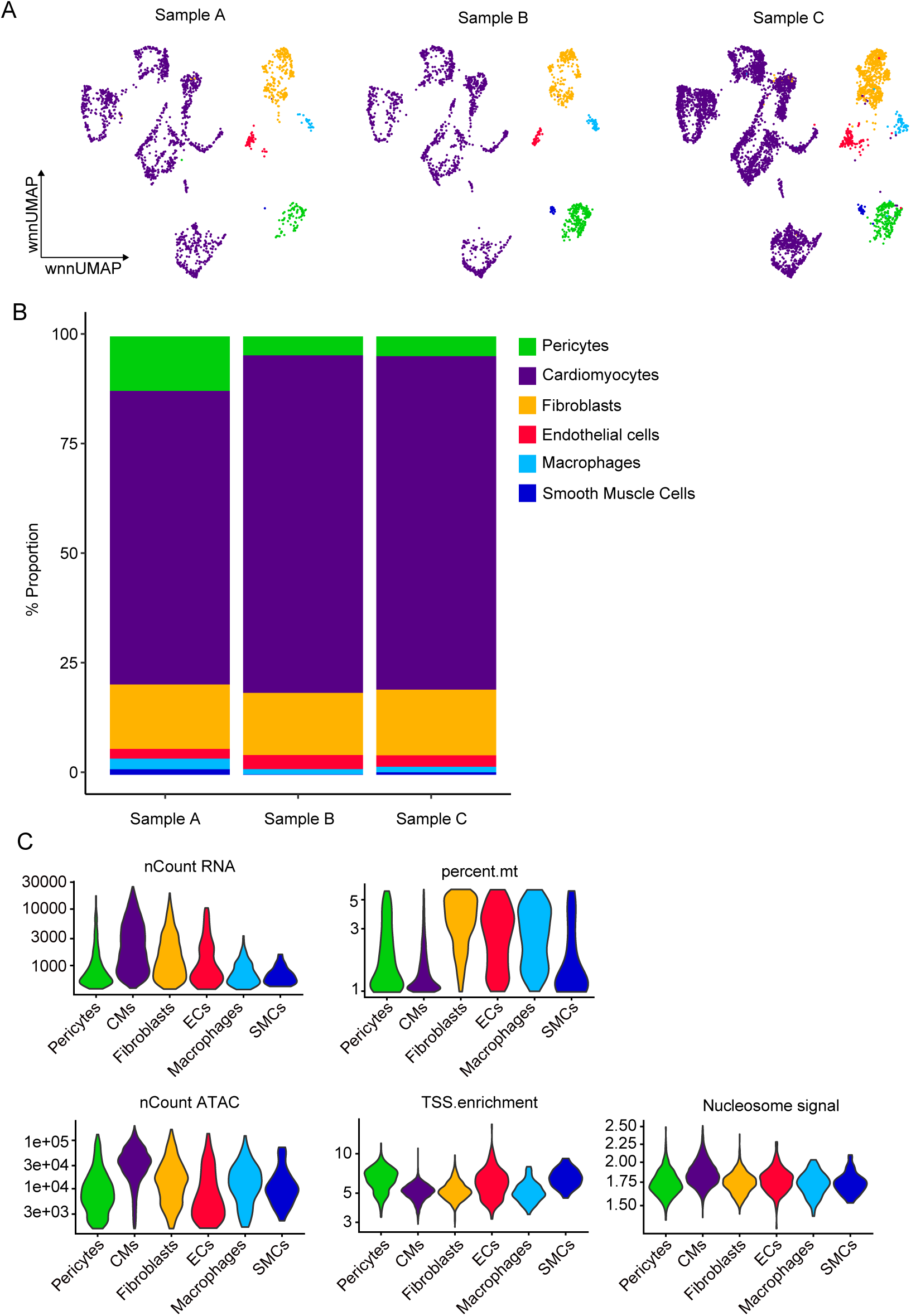
Composition of human cardiac cell types within human paired-multiomic dataset. (related to Figure 2). A. Weighted Nearest Neighbor (WNN) UMAPs of the three batches from which single-cell data was prepared. B. Stacked bar plot summarizing relative proportions of cell types from different batches. C. Data quality parameters for each major cardiac cell type.

**Supplementary Figure. 4.**
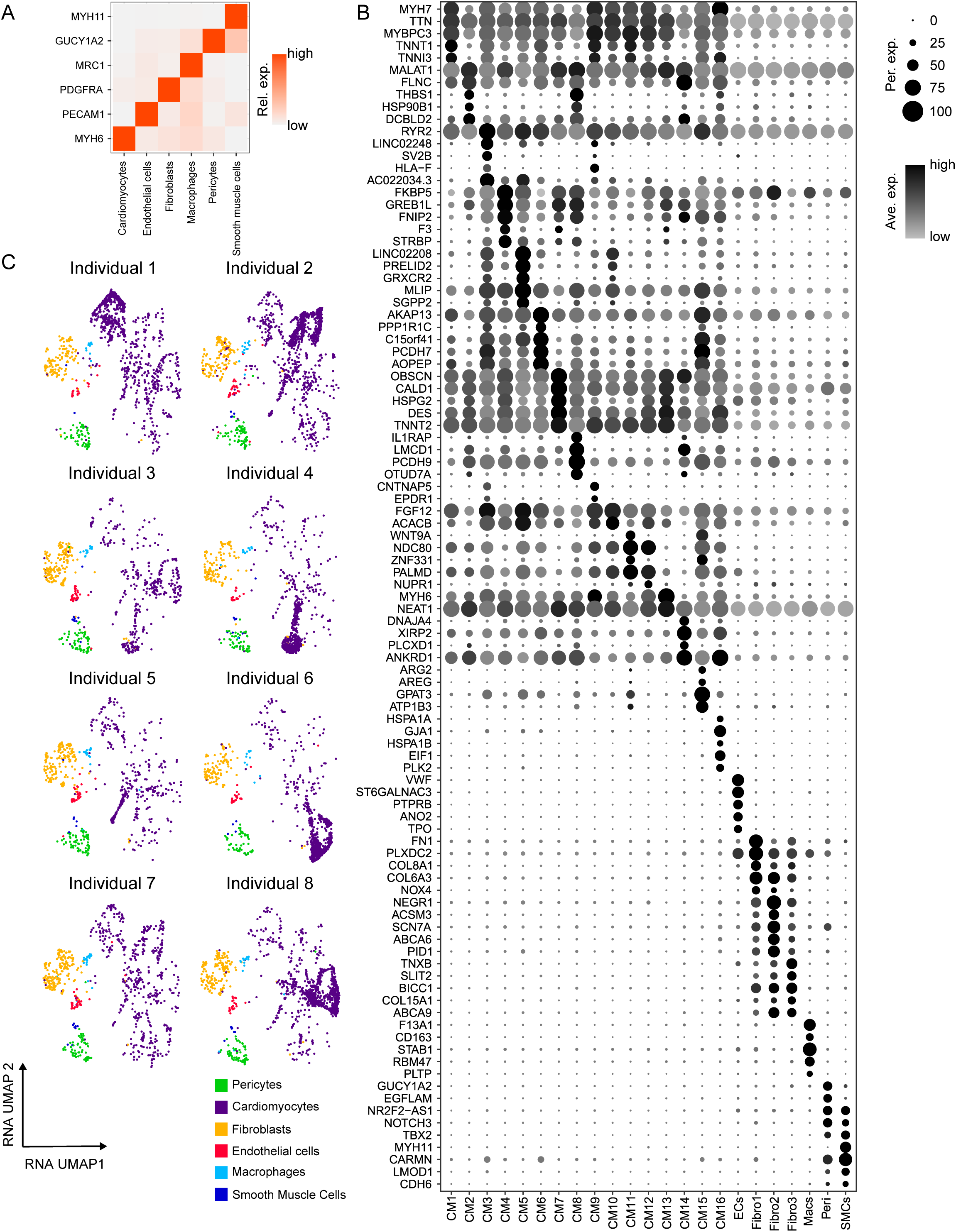
Markers of cardiac cell types within the human paired-multiomic dataset and sample deconvolution. A. Heat map of relative expression of canonical cell marker genes of major human cardiac cell types. B. Dot plot of the relative expression of the top 5 marker genes per cell cluster. Dot size and intensity indicate the proportion of cells expressing the genes relative average expression level, respectively, within each cell population. C. UMAP of cells from individual donor samples following deconvolution (see Methods).

**Supplementary Figure 5.**
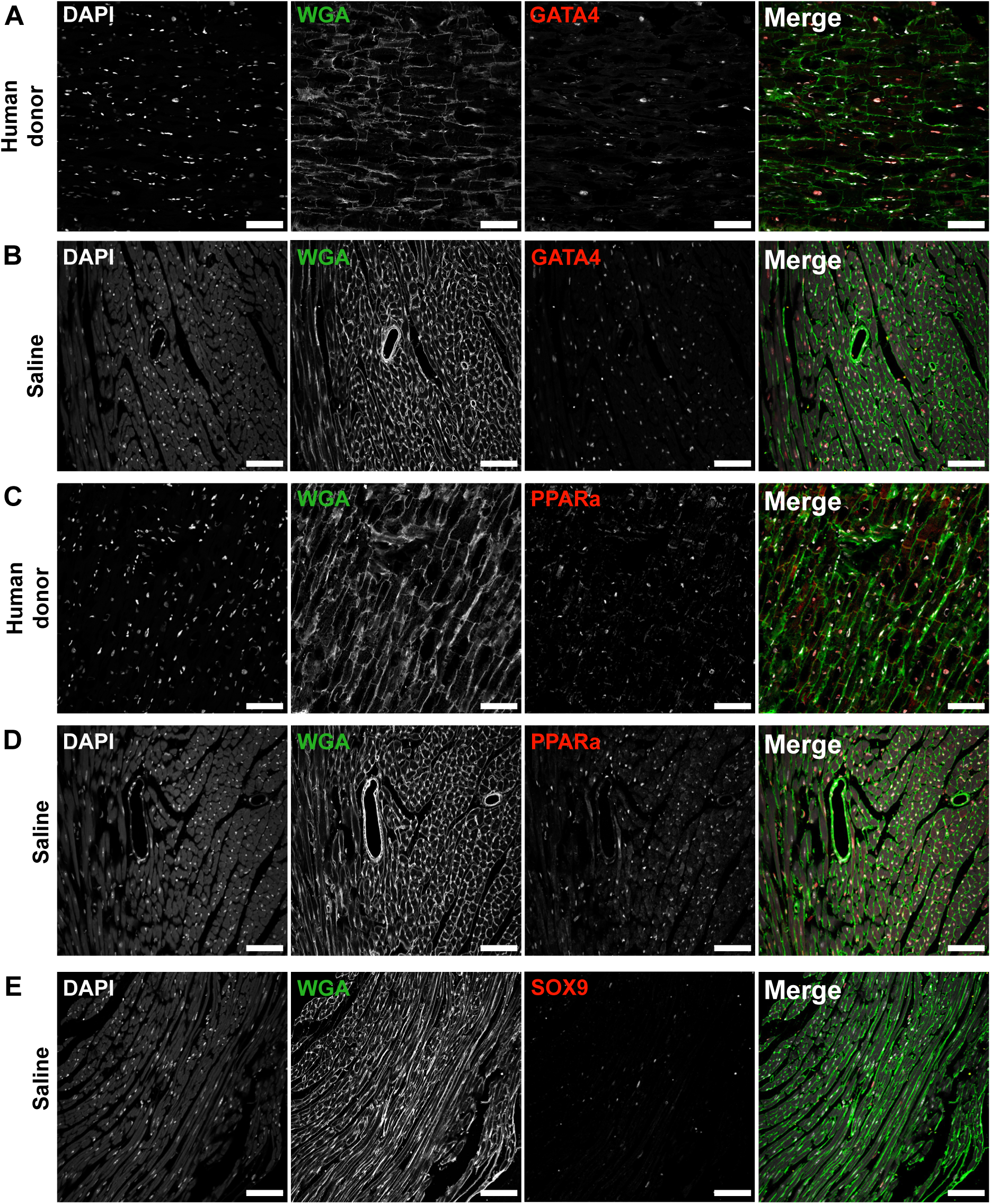
Expression of key cardiac transcription factors in mouse and human hearts. A. GATA4+ nuclei in a donor heart. B. GATA4+ nuclei in unstressed mouse cardiac tissue. C. PPARα+ nuclei in a donor heart. D. PPARα+ nuclei in an unstressed mouse cardiac tissue. E. SOX9+ nuclei in an unstressed mouse heart. Scale bars indicates 100 µm. DAPI, 4′,6-diamidino-2-phenylindole; WGA, wheat germ agglutinin.

**Supplementary Figure 6.**
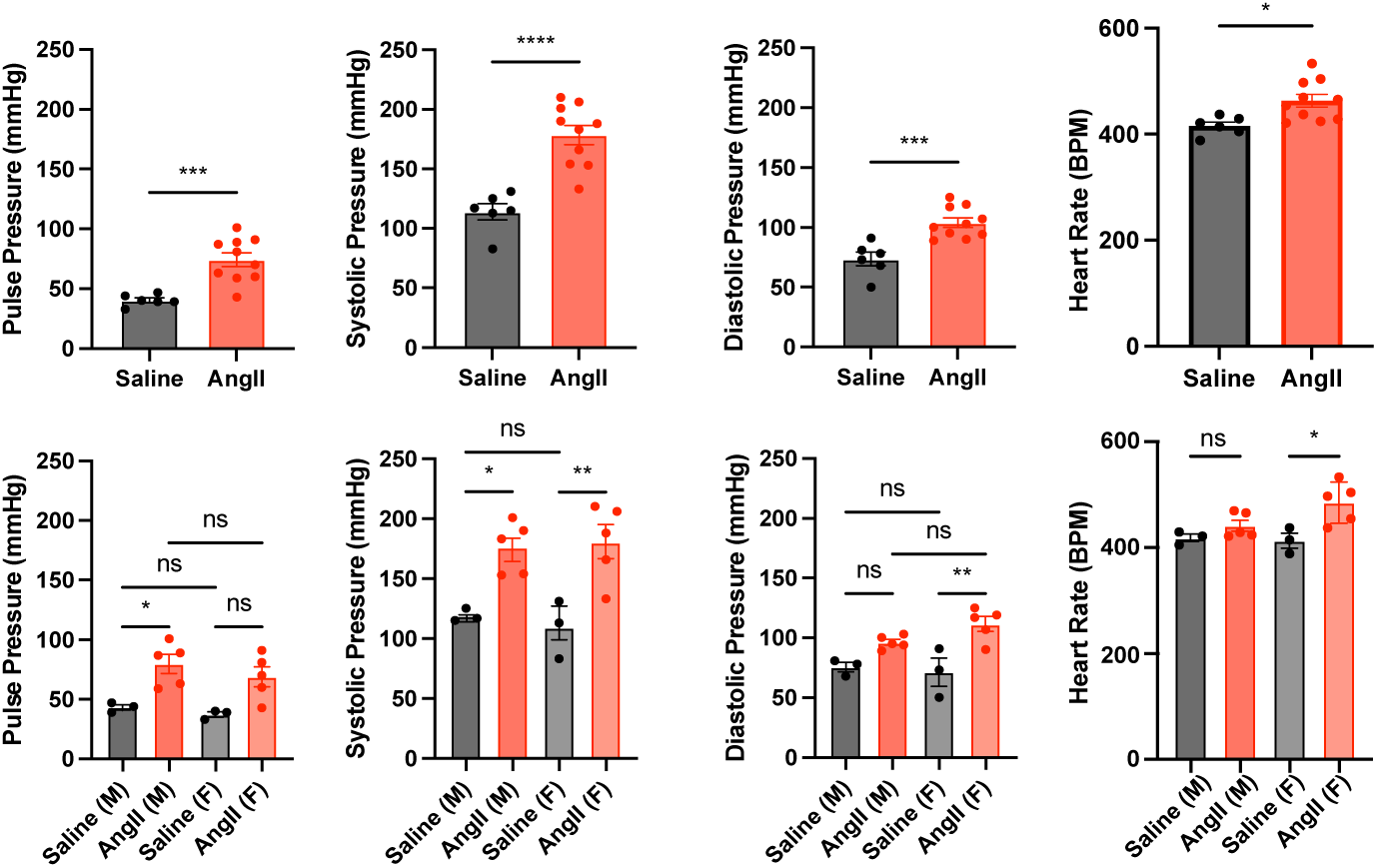
Assessment of hemodynamics and cardiac function after AngII infusion.

**Supplementary Figure 7.**
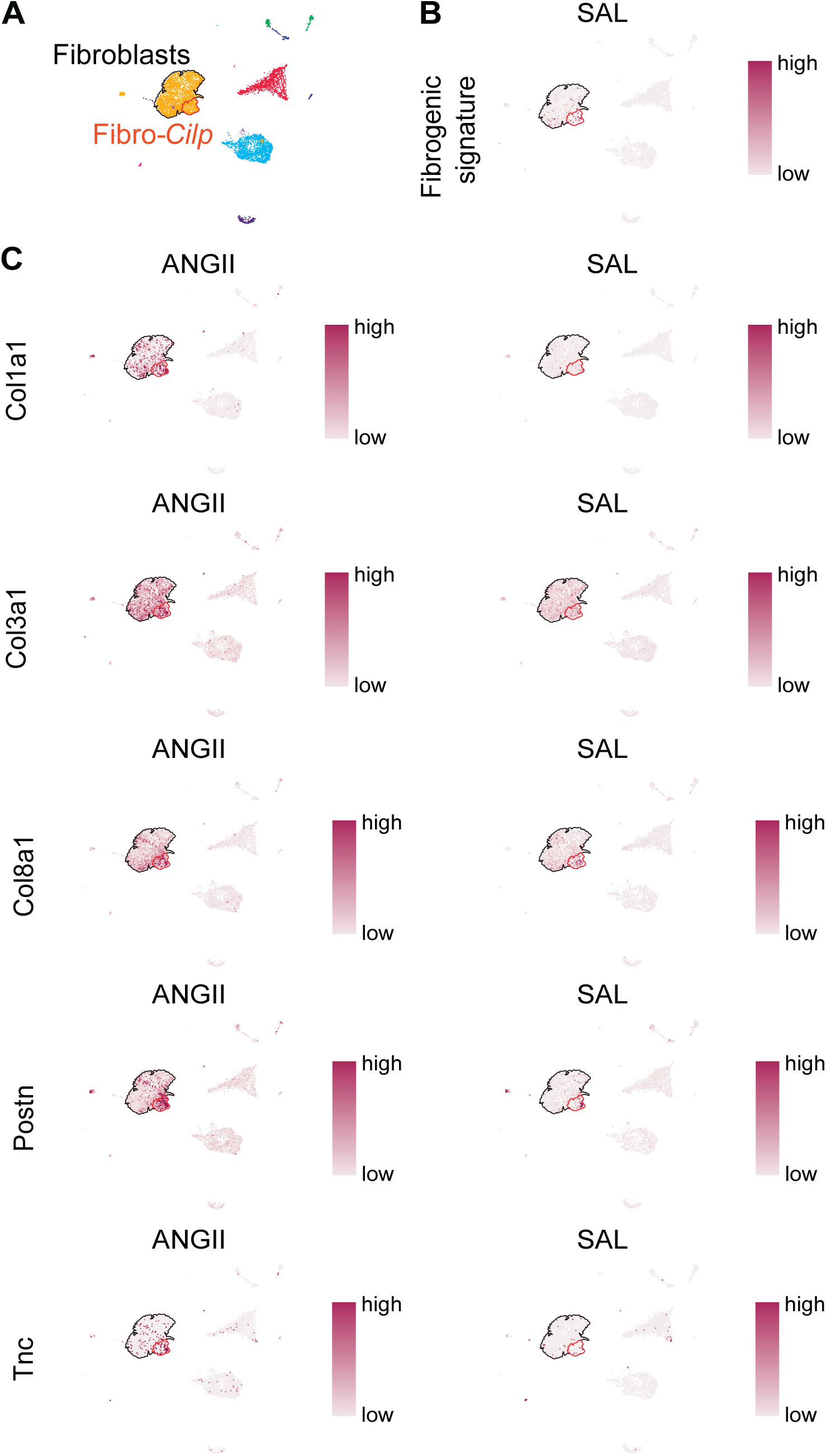
‘Fibrogenic signature’ module gene expression in AngII model (related to Figure 4B). A. UMAP of cardiac cell populations with Fibro-Cilp population and other fibroblasts highlighted. B. UMAP showing summed average gene expression corresponding to fibrogenic module for cells from Saline mouse hearts. C. UMAPs showing expression of individual genes comprising fibrogenic module in cells from AngII and Saline mouse groups.

**Supplementary Figure 8.**
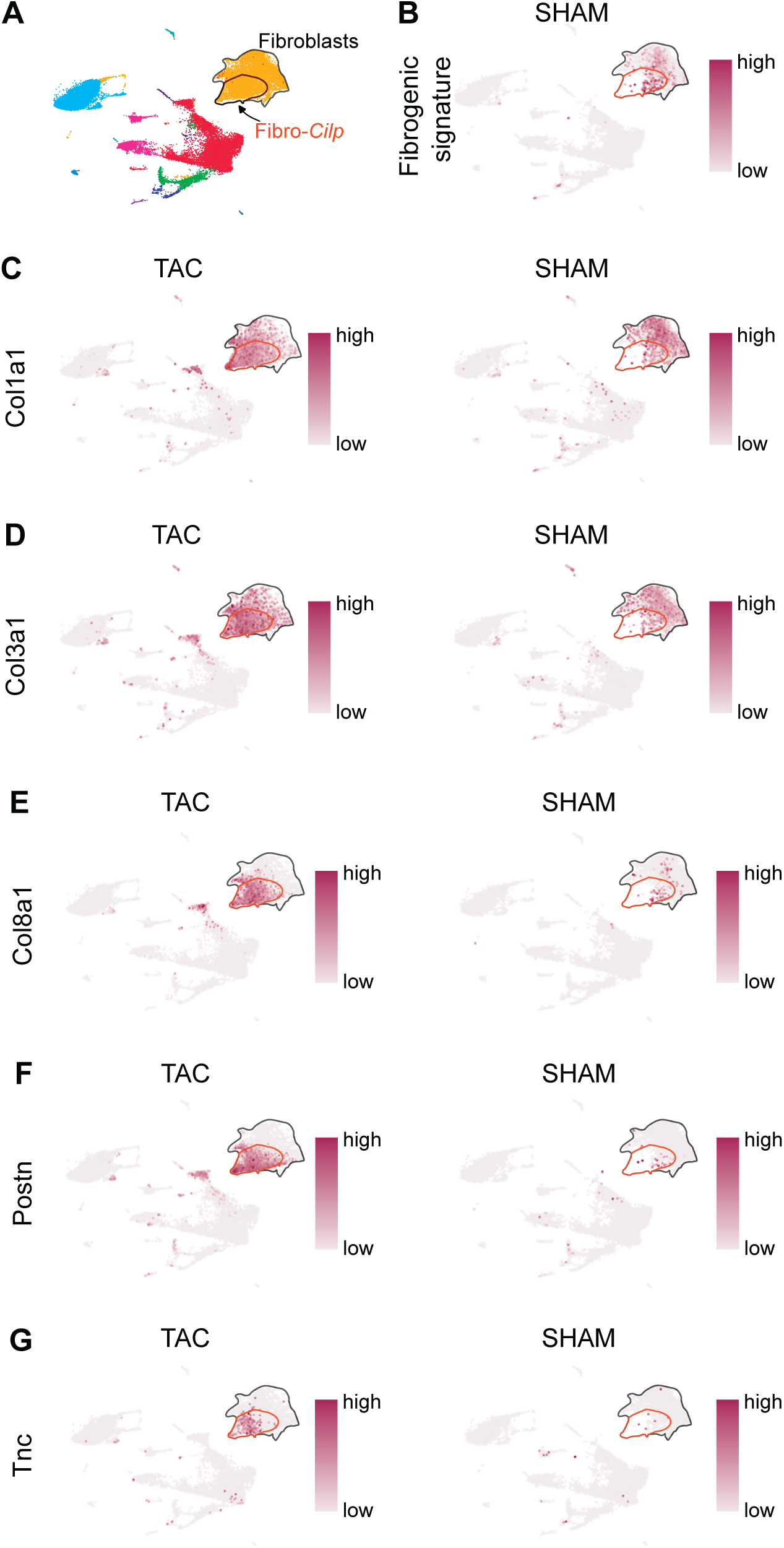
‘Fibrogenic signature’ module gene expression in TAC model (related to Figure 4E). A. UMAP of cardiac cell populations with Fibro-Cilp population and other fibroblasts highlighted. B. UMAP showing summed average gene expression corresponding to fibrogenic module for cells from Sham mouse hearts. C. UMAPs showing expression of individual genes comprising fibrogenic module in cells from TAC and Sham mouse groups.

**Supplementary Figure 9.**
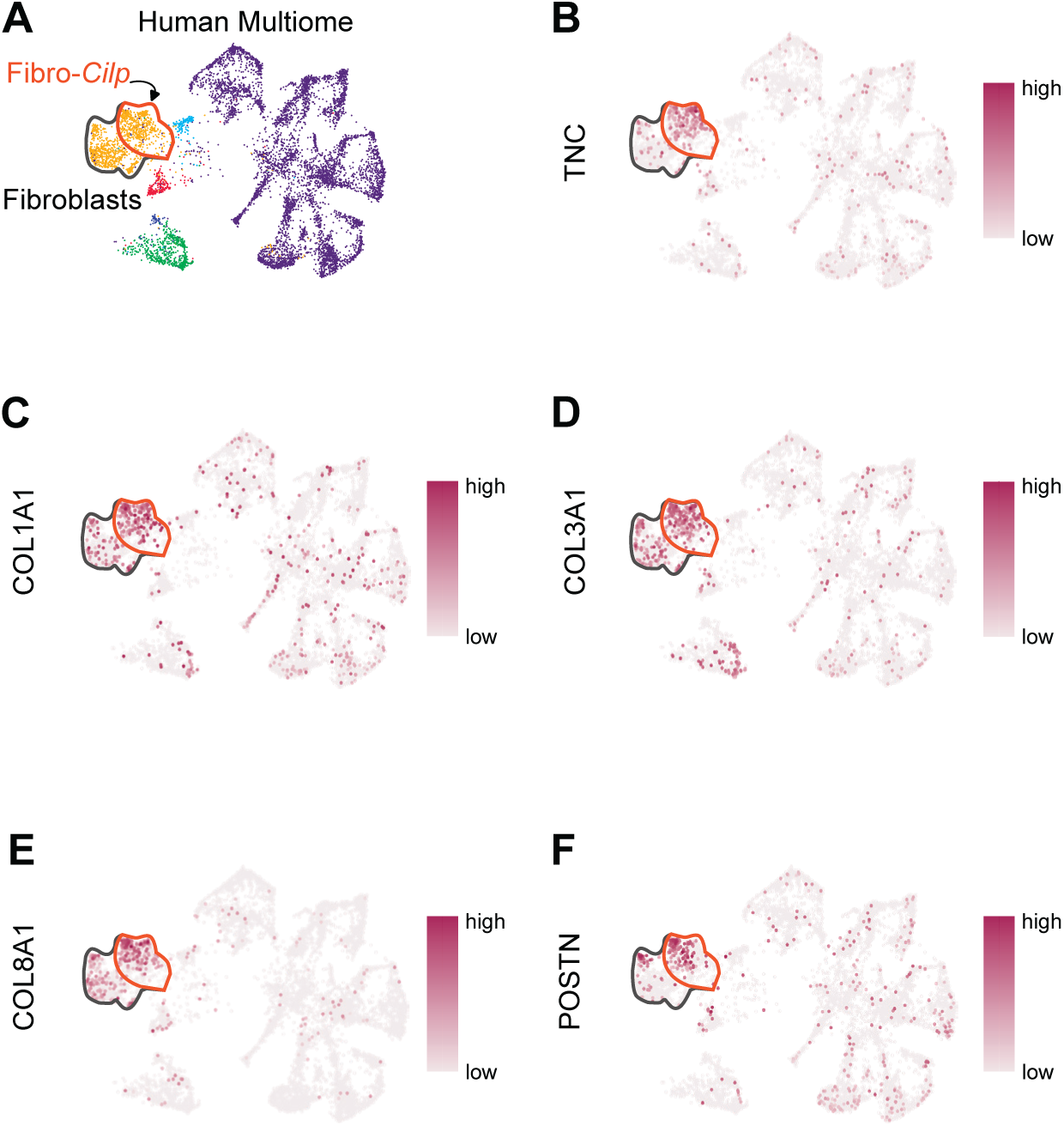
‘Fibrogenic signature’ module gene expression in human donor heart cells (related to Figure 4H). A. UMAP of cardiac cell populations with Fibro-Cilp population and other fibroblasts highlighted. **B-D**. UMAPs showing expression of individual genes comprising fibrogenic module in cells from TAC and Sham mouse groups.

**Supplementary Figure 10.**
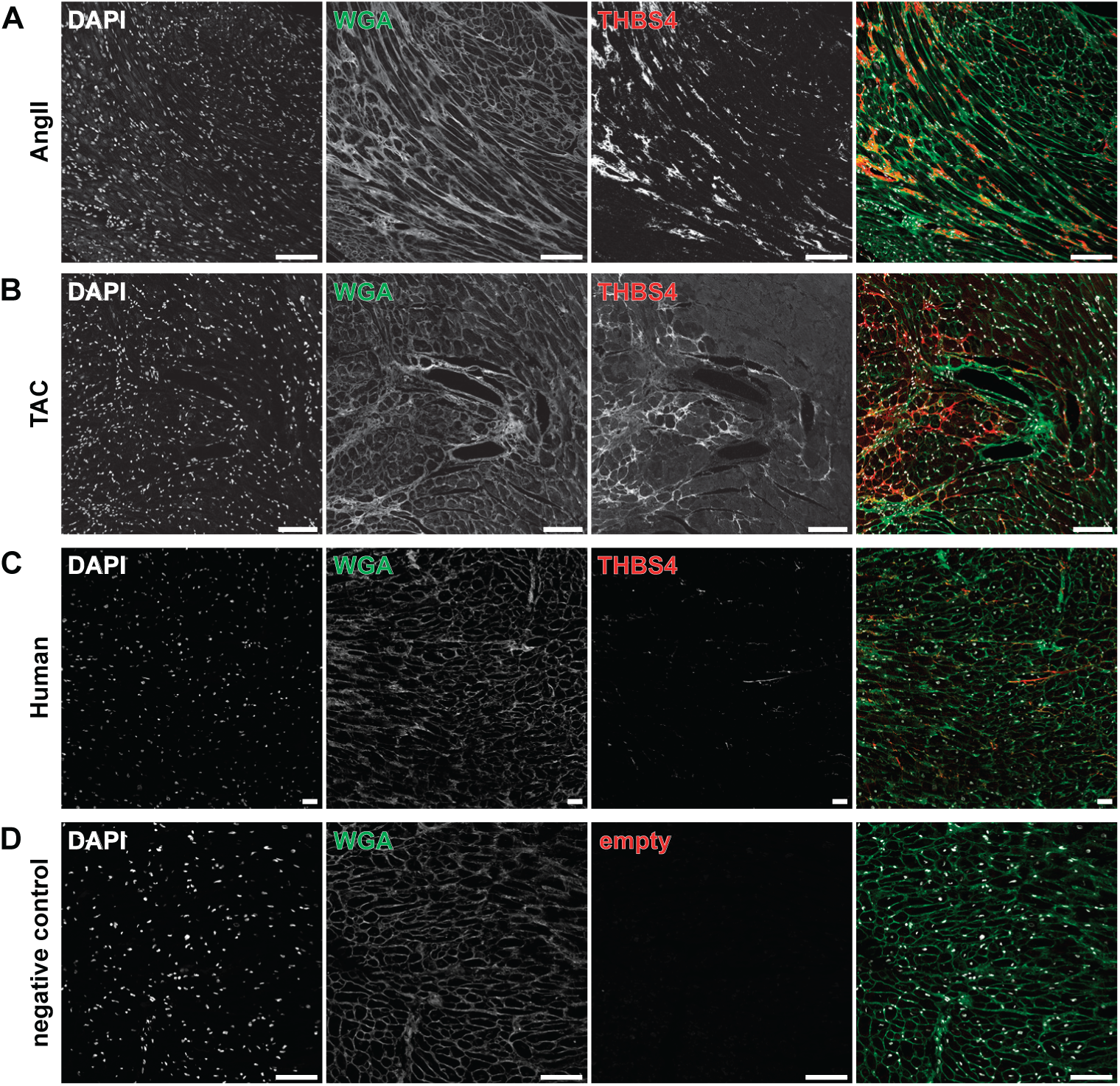
Validation of Thbs4 expression in different contexts. A. THSB4+ nuclei in an AngII mouse cardiac tissue. B. THSB4+ nuclei in a TAC mouse cardiac tissue. C. THSB4+ nuclei in human donor cardiac tissue. D. THSB4+ nuclei in an unstressed mouse cardiac tissue. Scale bar indicates 100 µm. DAPI indicates 4′,6-diamidino-2-phenylindole; WGA indicates wheat germ agglutinin.

**Supplementary Figure 11.**
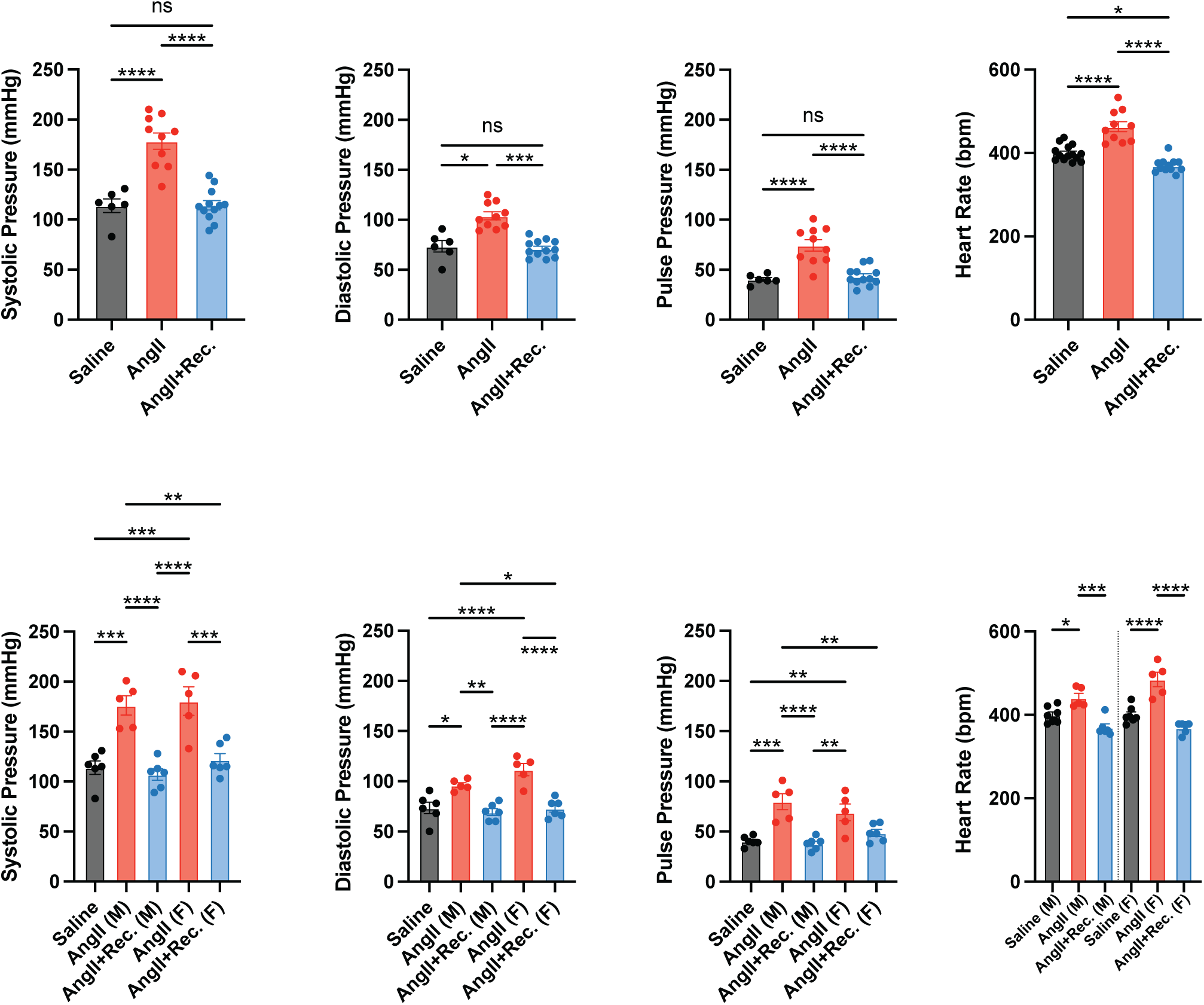
Assessment of physiological parameter in AngII and AngII+Rec mice.

**Supplementary Figure 12.**
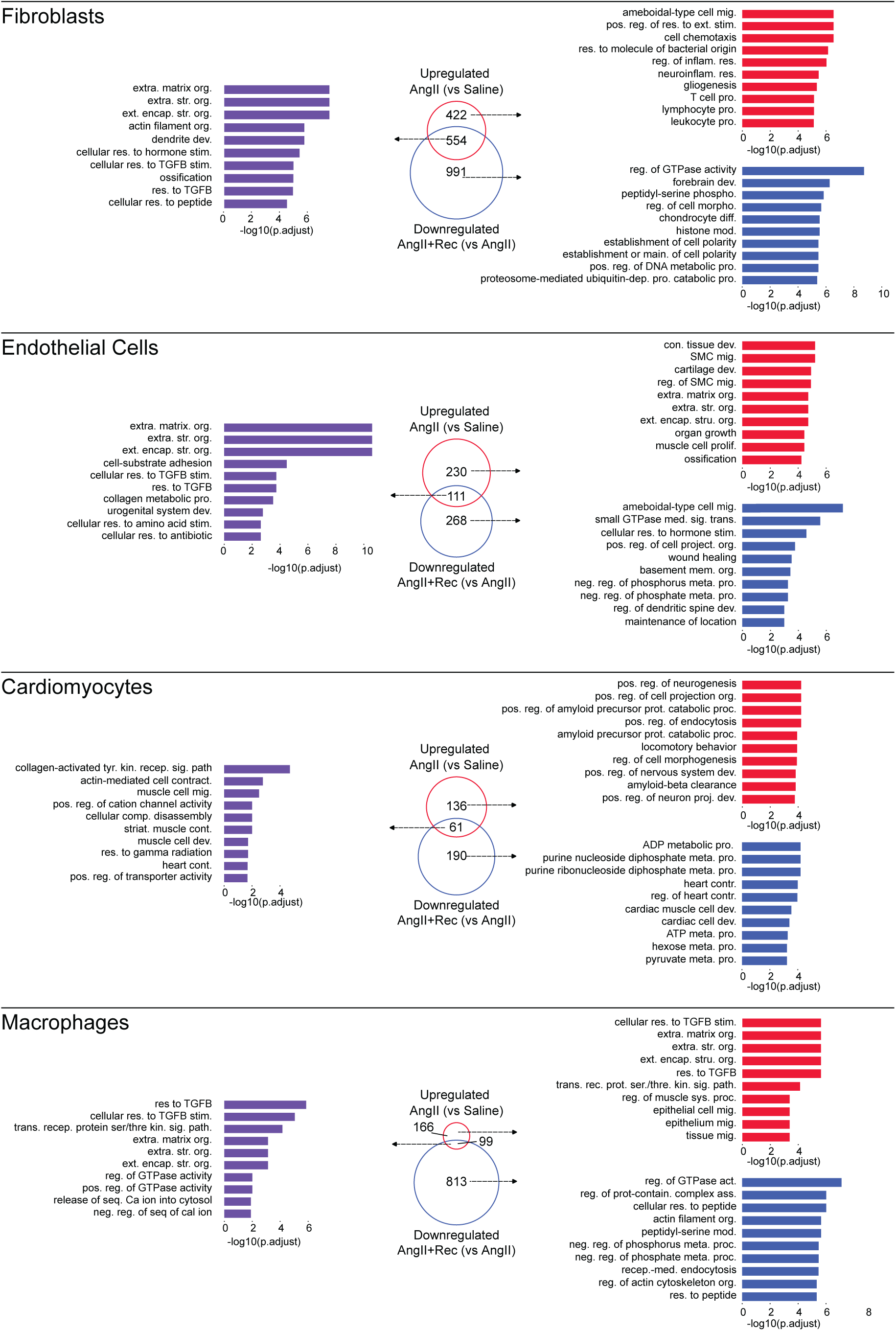
Assessment of cell population-specific gene programs reversibly induced by mouse AngII infusion. Venn diagrams summarize genes within major cell populations upregulated in AngII (vs Saline) mouse hearts and downregulated in AngII+Rec (vs AngII) mouse hearts for each major cardiac cell population. Purple bar plots represent top 10 enriched GO terms for genes that are up- and downregulated in AngII vs Saline and AngII+Rec vs AngII. Red and blue bar plots represent top 10 GO terms corresponding to genes that are uniquely up- or downregulated in AngII vs Saline, or AngII+Rec vs AngII, respectively.

**Supplementary Figure 13.**
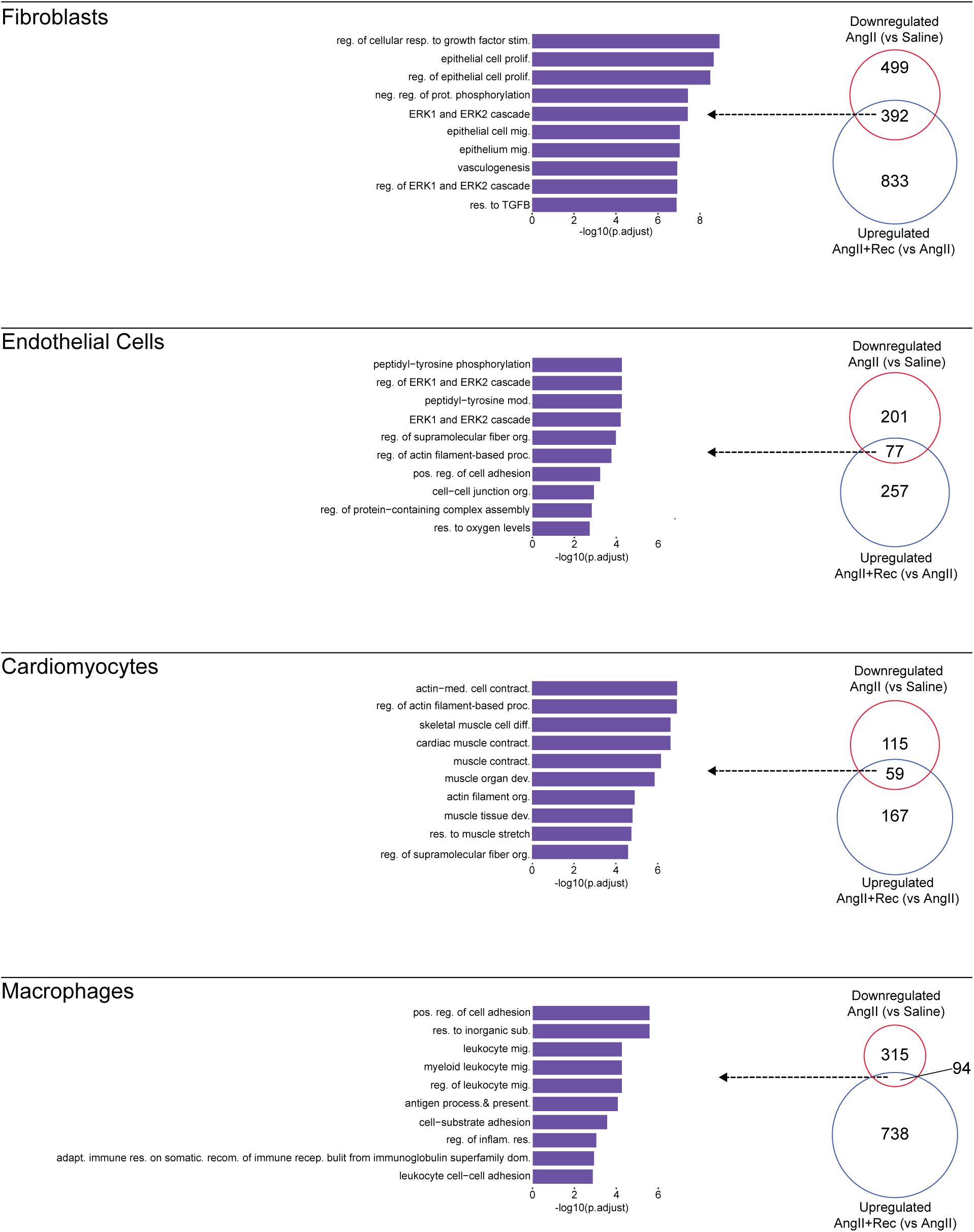
Assessment of cell population-specific gene programs reversibly suppressed by mouse AngII infusion. Venn diagrams summarize genes within cell populations downregulated in AngII (vs Saline) mouse hearts and upregulated in AngII+Rec (vs AngII) mouse hearts for each major cardiac cell population. Purple bar plots represent top 10 enriched GO terms for genes that are down- and upregulated in AngII vs Saline and AngII+Rec vs AngII.As in A but for macrophages.

**Supplementary Figure 14.**
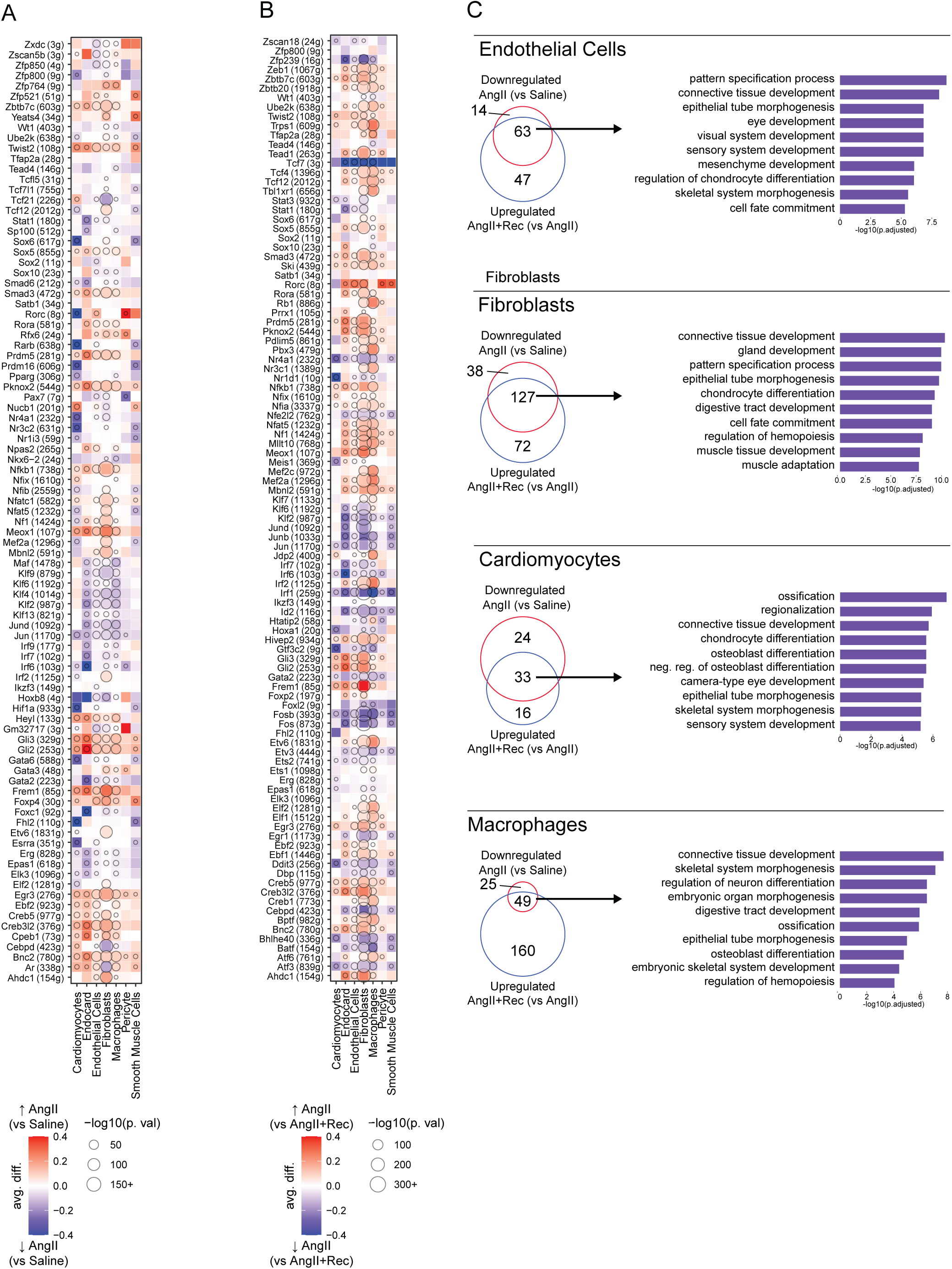
Reversibility of AngII-induced TF activity. A. Dot plot showing the eRegulon enrichment change on color scale and p-value significance in circle size, comparing major cell types within AngII to Saline mouse hearts. Red color indicates eRegulons that have increased in enrichment, and blue those that have decreased, in AngII mouse heart cells compared to those from Saline. B. Dot plot showing the eRegulon enrichment change on color scale and p-value significance in circle size, comparing major cell types within AngII to AngII+Rec mouse hearts. Red color indicates eRegulons that have increased in enrichment, and blue those that have decreased, in AngII mouse heart cells compared to those from AngII+Rec. C. Venn diagram showing overlap of eRegulons increased in enrichment in AngII mouse hearts compared to Saline mouse hearts, and eRegulons that are enriched in AngII mouse hearts compared to AngII+Rec mouse hearts. GO enrichment analysis (purple bars) was performed on a overlapping eRegulons from these two lists for each major cardiac cell type.

**Supplementary Figure 15.**
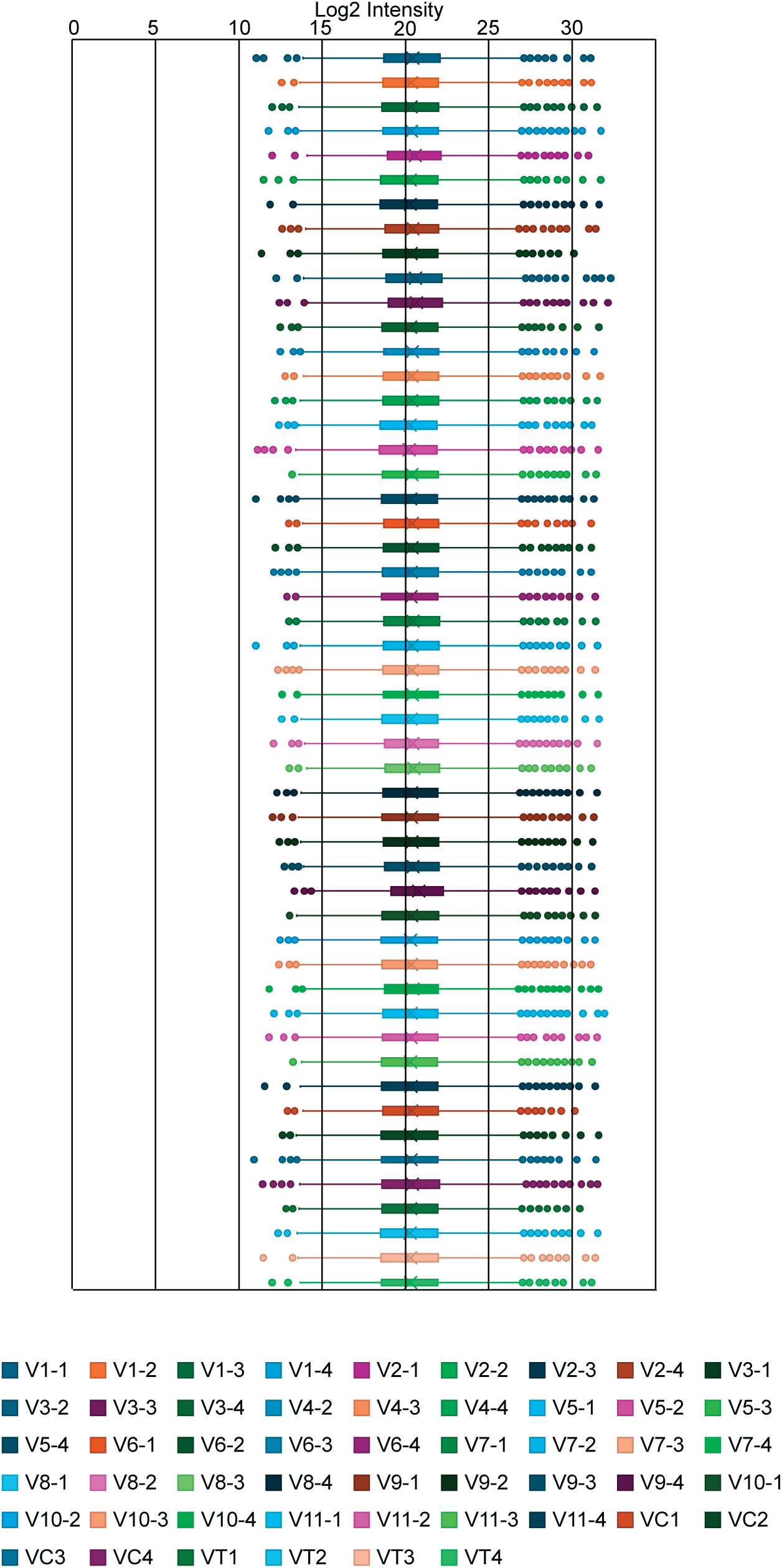
Proteomic normalization of human ventricular fibroblast samples. Distribution of protein abundances of cardiac fibroblasts samples.

## Notes

### Competing Interest Statement

The authors have declared no competing interest.

### Summary of Updates

Minor amendment to some figure legends and clarity of some supplementary figures.

